# PDLP5 regulates aquaporin-mediated hydrogen peroxide transport in *Arabidopsis*

**DOI:** 10.64898/2026.01.21.700913

**Authors:** Zhongpeng Li, Su-Ling Liu, Saiful Islam, Madelyn Clements, Yani Chen, Kyaw Aung

## Abstract

Aquaporins facilitate the diffusion of water and small neutral molecules, including hydrogen peroxide (H_2_O_2_). In *Arabidopsis*, plasma membrane intrinsic proteins (PIPs, members of aquaporins) mediate H_2_O_2_ transport; however, the regulatory mechanisms governing their activity are not fully understood. Here, we show that plasmodesmata-located protein 5 (PDLP5), a key regulator of plasmodesmal function, negatively regulates PIP-dependent H_2_O_2_ transport. PDLP5 forms complexes with PIPs, with its extracellular domain predicted to associate with the extracellular face of the channel. PDLP5 overexpression suppresses H_2_O_2_ uptake and confers reduced sensitivity to H_2_O_2_-induced root growth inhibition, whereas *pdlp5* mutants display the opposite phenotype. We further identify PIP2;5 as a critical component of PDLP5-mediated regulation. These findings reveal an extracellular mechanism for modulating aquaporin function, expanding current models of channel regulation in plants.

## Introduction

Reactive oxygen species (ROS) play a central role in plant development and environmental responses. Although potentially damaging, ROS act as versatile signaling molecules that regulate a broad spectrum of cellular processes, including growth, differentiation, and immunity^1,2^. Among the major ROS species, hydrogen peroxide (H_2_O_2_) is relatively stable and diffuses over cellular and tissue distances, making it a key redox signal. In plants, H_2_O_2_ is produced in the apoplast and within organelles, including chloroplasts, mitochondria, and peroxisomes^3,4^. Maintaining H_2_O_2_ homeostasis requires precise coordination of its production, scavenging, and transport across membranes.

Aquaporins (AQPs) are integral membrane channels that facilitate the diffusion of water and small neutral solutes, including H O ^5^. In plants, AQPs are classified into four major intrinsic protein subfamilies: plasma membrane intrinsic proteins (PIPs), tonoplast intrinsic proteins (TIPs), nodulin-26-like intrinsic proteins (NIPs), and small basic intrinsic proteins (SIPs)^6^. The Arabidopsis genome encodes 13 PIPs, grouped into two subfamilies, PIP1 and PIP2^7,8^. Several PIPs, including PIP1;4 and PIP2;1, mediate H_2_O_2_ transport and are essential for immune signaling and stress tolerance^9–12^. During pattern-triggered immunity, apoplastic ROS produced by the plasma membrane (PM)-localized NADPH oxidases respiratory burst oxidase homologue D (RBOHD) and RBOHF are proposed to enter the cytosol through PIPs, thereby activating intracellular redox signaling cascades^13,14^. However, the regulatory mechanisms that modulate PIP-dependent H_2_O_2_ transport remain poorly understood.

Emerging evidence suggests that ROS signaling is closely coordinated with the regulation of plasmodesmata, which are membrane-lined conduits connecting adjacent cells. The plasmodesmal PM (PD-PM), plasmodesmal endoplasmic reticulum (PD-ER), and cytoplasmic sleeve form a dynamic conduit. The plasmodesmata-dependent permeability is controlled by the deposition of callose (β-1,3-glucan)^15,16^. Callose accumulation narrows the plasmodesmal aperture and restricts intercellular transport, whereas its degradation reopens the channel^17^. In addition, multiple C2 domains transmembrane domain proteins (MCTPs) are targeted to PD-ER and function as tethers between PD-ER and PD-PM, regulating plasmodesmal function in a callose-independent manner^18^.

Plasmodesmata-located protein 5 (PDLP5) is preferentially localized at PD-PM and functions as a key regulator of plasmodesmal aperture^19–22^. PDLP5 contains two extracellular domains of unknown function 26, a transmembrane helix, and a short cytoplasmic tail^19^. Its expression is induced by salicylic acid and pathogen infection^20^. Both its induction and overexpression promote callose accumulation and restricted cell-to-cell movement of molecules^20,23^. Consistent with these functions, *pdlp5* mutants exhibit reduced callose deposition and enhanced plasmodesmal permeability. PDLP5 is also required for systemic acquired resistance^24^ and for RBOHD-dependent plasmodesmal closure through a PDLP1-PDLP5-NHL3 (non-race-specific disease resistance/hin1 hairpin-induced-like protein 3) complex^22^. Given its central role in regulating plasmodesmal function, PDLP5 contributes broadly to plant growth, development, and immunity^21,23,25–27^. Despite its involvement in diverse cellular and physiological processes, the molecular function of PDLP5 remains unknown.

Here, we demonstrate that PDLP5 physically interacts with PIPs and inhibits PIP2;5-mediated H_2_O_2_ transport. This reveals a previously unrecognized mechanism by which a plasmodesmal protein modulates aquaporin function, linking the regulation of intercellular connectivity to the control of cellular redox signaling.

## Results

### PDLP5 physically interacts with PIPs

To investigate the molecular function of PDLP5, we examined how it operates in concert with its interacting proteins. Previously, we performed a proximity labeling assay to identify potential interacting proteins of PDLP5^23^. Compared with MCTP3, PDLP5 significantly enriched 9 out of 13 PIP members^23^, suggesting a potential interaction between PDLP5 and PIPs. To test for direct physical interactions, we first performed split-ubiquitin yeast two-hybrid (suY2H) assays. Co-expression of PDLP5-Cub with NubWT and NubG served as positive and negative controls, respectively. The positive control showed blue coloration on X-Gal medium and strong growth on stringent medium containing 3-amino-1,2,4-triazole (3-AT) or methionine (Met), whereas the negative control remained white and exhibited weak growth (Fig. 1a). Co-transformation of PDLP5-Cub with NubG-PIPs resulted in blue coloration on X-Gal medium and enhanced growth on stringent media, indicating an interaction between PDLP5 and PIPs (Fig. 1a).

**Fig. 1:**
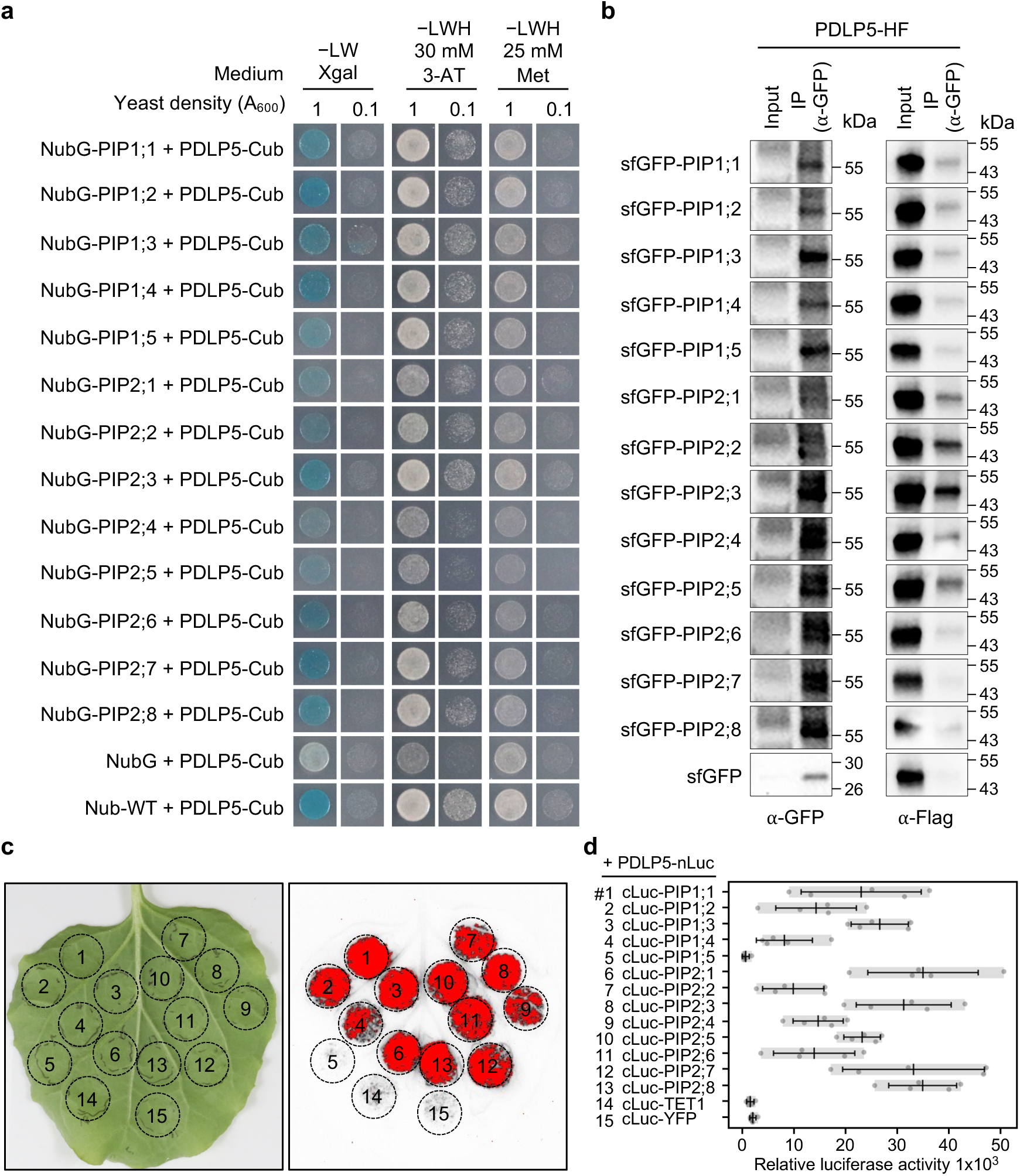
PDLP5 physically interacts with PIPs. **a**, Split-ubiquitin yeast two-hybrid assays determine the interaction between PDLP5 and PIPs. Nub-WT and NubG were used as positive and negative controls, respectively. To conduct blue/white screening, −LW + X-Gal synthetic dropout plates were utilized, while −LWH synthetic dropout plates were used for histidine auxotrophy selection. To increase the stringency of the assay, 30 mM 3-AT or 25 mM Met was added to the −LWH synthetic dropout plates. **b**, Co-immunoprecipitation assays indicate the presence of protein complexes containing PDLP5 and PIPs. Agrobacteria harboring different plasmids were co-infiltrated into *N. benthamiana*. Samples were collected 2 days after infiltration and subjected to co-IP. A GFP antibody was used to detect the expression of sfGFP fusion protein and the enrichment of sfGFP fusion proteins by GFP-Trap. A Flag antibody was used to detect the expression of HF fusion protein and the interaction between HF and sfGFP fusion proteins. A free sfGFP fusion protein was included as a negative control. **c,** Luciferase complementation assays demonstrate the interaction between PDLP5 and PIPs. Agrobacteria carrying the respective plasmids were co-infiltrated into *N. benthamiana* leaves. Two days after infiltration, leaves were sprayed with 1 mM D-luciferin, and luminescence was imaged using an iBright system (left). **d**, Quantification of luminescence was performed using a microplate reader (right). Data represent the mean ± SD (n = 5).

We next verified the interaction between PDLP5 and PIPs *in planta* using co-immunoprecipitation (co-IP) assays. sfGFP-PIPs and PDLP5-HF were transiently co-expressed in *Nicotiana benthamiana*. As shown in Fig. 1b, all sfGFP-PIPs, with the exception of sfGFP-PIP2;7, co-immunoprecipitated PDLP5-HF, whereas free sfGFP did not. The results indicate that PDLP5 and PIPs associate within the same protein complex *in planta*. To further confirm these interactions, we performed a luciferase complementation assay in *N. benthamiana*. Co-expression of PDLP5-nLuc with cLuc-PIPs produced strong luminescence signals (except cLuc-PIP1;5), whereas co-expression with cLuc-tetraspanin1 (TET1, a PM-associated protein^28^) or cLuc-YFP did not (Fig. 1c). Luminescence intensity measurements supported these observations, with significantly higher signals detected in samples co-expressing cLuc-PIPs (except cLuc-PIP1;5) compared to negative controls (Fig. 1d). Together, these results show that PDLP5 physically interacts with PIP proteins.

### AlphaFold predicts physical interactions between PDLP5 and AQPs

We next utilized AlphaFold-based predictions to model PDLP5-PIP complexes. We reasoned that these analyses could be informed by the availability of experimentally determined PDLP5 ecodomain structures, which were solved at a resolution of 1.25 Å by X-ray crystallography^29^. In addition, high-resolution crystal structures for aquaporins are available^30–33^. We used AlphaFold3^34^ to model PDLP5-PIP heterodimers. Extended Data Fig. 1a shows the formation of a PDLP5-PIP1;1 complex, in which the PDLP5 ectodomain interacts with the extracellular face of the PIP1;1 channel. Because AQPs function as tetramers^35^, we next used AlphaFold3 to predict whether PDLP5 could associate with PIP tetramers. The analysis predicted an octameric PDLP5-PIP1;1 complex comprising four PDLP5 and four PIP1;1 subunits (Extended Data Fig. 1b). In this model, the PDLP5 ectodomain is positioned at the extracellular channel opening of PIP1;1, forming a tightly fitted complex in which PDLP5 appears to occlude the PIP pores. The predicted PDLP5-PIP interfaces are supported by relatively low interfacial predicted TM (ipTM) scores (Extended Data Fig. 1), a metric provided by AlphaFold3 that estimates the confidence in intermolecular interfaces. Nevertheless, multiple independent experimental observations provide strong empirical support for a physical interaction between PDLP5 and PIPs.

Given that AQPs are highly conserved across species, from archaea to humans^36^, we also used AlphaFold3 to model complexes between PDLP5 and human aquaporins (HsAQPs). PDLP5 was predicted to form both heterodimeric and hetero-octameric complexes with HsAQPs. Extended Data Fig. 1c shows the predicted PDLP5-HsAQP0 heterodimer. Consistent with these structural predictions, suY2H and luciferase complementation assays revealed positive interactions between PDLP5 and several HsAQPs (Extended Data Fig. 2). Together, the structural models and interaction assays suggest that the PDLP5 ectodomain can associate with the extracellular channel openings of HsAQPs, indicating a conserved structural compatibility between PDLP5 and AQPs across kingdoms.

### PDLP5 negatively regulates H_2_O_2_ uptake

AQPs facilitate the transport of water, H_2_O_2_, and other small neutral molecules^5^. Because the ectodomain of PDLP5 is predicted to interact with the extracellular face of PIP channels, we hypothesized that PDLP5 modulates PIP-mediated transport of small molecules across the PM. We focused on H_2_O_2_ for the following reasons: (1) PDLP5 plays a critical role in bacterial immunity^20,27^; (2) bacterial infection and perception of pathogen-associated molecular patterns, such as the 22-amino acid peptide flg22 derived from bacterial flagellin, trigger a rapid ROS burst through RBOHD and RBOHF^37^; (3) PDLP5 is required for systemic ROS signaling during light stress^38^; and (4) proximity labeling identified multiple ROS-related proteins, including those involved in ROS production, sensing, and transport, as putative PDLP5 interactors^23^. Consistent with these observations, several Arabidopsis PIPs have been shown to facilitate H_2_O_2_ transport^9,10,39,40^.

We first examined whether PDLP5 influences ROS production during the flg22-induced ROS burst. As shown in Figs. 2a and 2b, *pdlp5* knockout mutant and *PDLP5-HF* overexpressor exhibited comparable levels of flg22-induced ROS with Col-0, indicating that PDLP5 does not substantially affect flg22-triggered ROS production. Next, we investigated whether PDLP5 modulates the transport of ROS. If PDLP5 restricts H_2_O_2_ transport, differences in extracellular ROS (eROS) accumulation should be detected among Col-0, *pdlp5*, and *PDLP5-HF* plants. To assess this, leaves were treated with flg22 to induce eROS production, and eROS accumulation was detected using Amplex UltraRed (AUR), a membrane-impermeable probe that reacts with H_2_O_2_ to yield a fluorescent product^41^. As shown in Figs. 2c and 2d, *PDLP5-HF* plant accumulated significantly higher eROS compared with Col-0, whereas *pdlp5* mutant showed a reproducible but not statistically significant reduction. Because PDLP5 overexpression increased eROS accumulation without altering ROS production (Figs. 2a and 2b), these data suggest that PDLP5 restricts H_2_O_2_ transport from the apoplast into the cytosol. To further test this hypothesis, we detected intracellular ROS (iROS) using 2′,7′-dichlorodihydrofluorescein diacetate (H_2_DCFDA). After entering cells, H_2_DCFDA is hydrolyzed and oxidized by ROS, including H_2_O_2_, to yield fluorescent dichlorofluorescein (DCF)^42^. Following H_2_O_2_ treatment, protoplasts isolated from *pdlp5* leaves exhibited stronger DCF fluorescence compared with Col-0, whereas *PDLP5-HF* protoplasts showed markedly reduced fluorescence (Figs. 3a and 3b), indicating that PDLP5 suppresses H_2_O_2_ uptake. Similarly, *PDLP5-HF* root tips displayed significantly lower DCF fluorescence than Col-0 following H_2_O_2_ treatment (Figs. 3c and 3d). Together, these findings indicate that PDLP5 negatively regulates H_2_O_2_ transport into cells.

**Fig. 2:**
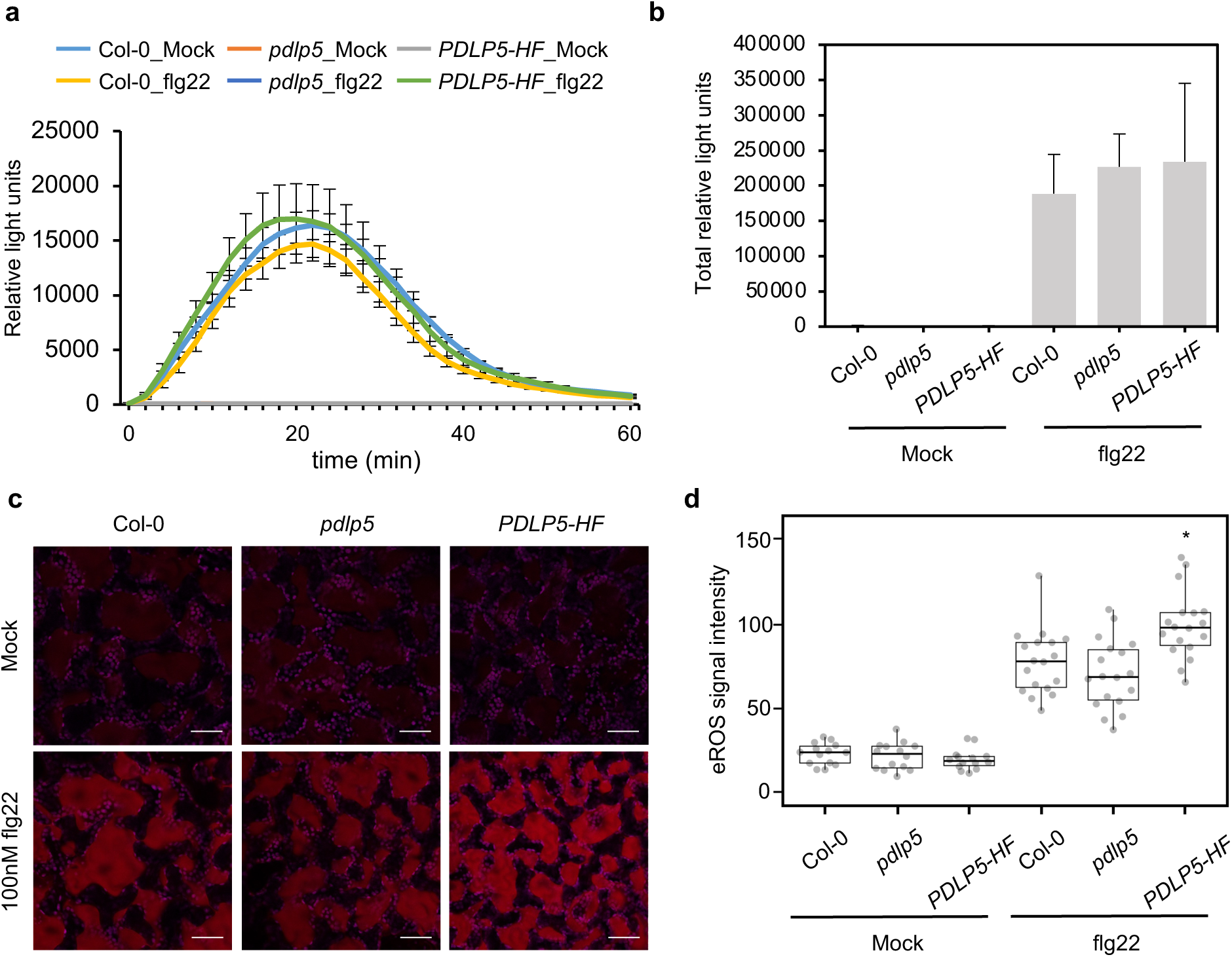
PDLP5 regulates eROS levels independently of the flg22-triggered oxidative burst. **a**, ROS production assay. Leaf discs from four-week-old Arabidopsis wild-type Col-0, *pdlp5* knock mutants, and *PDLP5-HF* overexpressors were examined for flg22-induced ROS burst with or without flg22 treatment. Error bars indicate the standard error of the mean (SEM); n = 8. **b,** Total ROS production measured over 60 minutes following with or without flg22 treatment. **c**, AUR staining of leaves upon flg22 treatment. Leaves from 3-week-old Arabidopsis were inoculated with 10 μM AUR and 100 nM flg22. Confocal images were taken 30 min after treatment. Red signals indicate AUR fluorescence, while magenta signals correspond to chlorophyll autofluorescence. Scale bars = 50 μm. **d,** Quantitative data show the flg22-induced accumulation of apoplastic H_2_O_2_. The box plot shows the mean with SD. n=14 (Mock) or 18 (flg22). Asterisks indicate statistically significant differences compared with wild-type Col-0 (*t*-Test; two-tailed; P < 0.05).

**Fig. 3:**
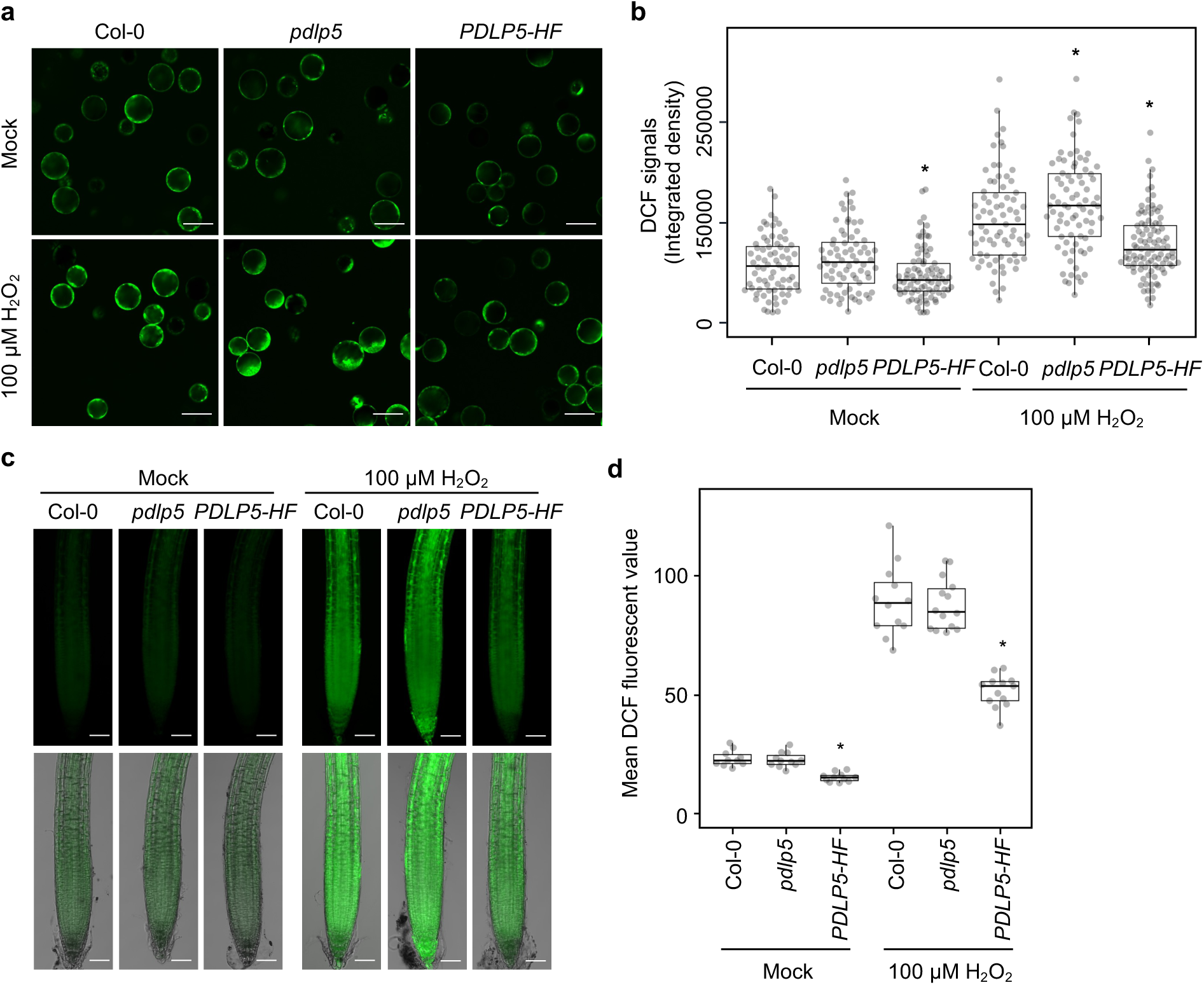
PDLP5 negatively regulates H_2_O_2_ transport in Arabidopsis. **a,** Confocal images show the uptake of H_2_O_2_ in Arabidopsis mesophyll protoplasts. Leaves from 4-week-old Arabidopsis were subjected to protoplast isolation. The protoplasts were incubated with 10 μM H_2_DCFDA and 100 μM H_2_O_2_. DCF signals were imaged 20 minutes after the treatment. Scale bars = 50 μm. **b,** Quantitative data show the accumulation of H_2_O_2_ in Arabidopsis mesophyll protoplasts. The box plot shows the mean with SD. Col-0: n = 72 (mock) and n = 73 (100 μM H_2_O_2_); *pdlp5*: n = 72 (mock) and n = 78 (100 μM H_2_O_2_); and *PDLP5-HF*: n = 83 (mock) and n = 97 (100 μM H_2_O_2_). Asterisks indicate statistically significant differences compared with wild-type Col-0 (*t*-Test; two-tailed; P < 0.05). **c,** Confocal images show the uptake of H_2_O_2_ in Arabidopsis root tips. 7-day-old Arabidopsis seedlings were incubated with 10 μM H_2_DCFDA for 15 minutes prior to H_2_O_2_ treatment. DCF signals were imaged 10 minutes after the H_2_O_2_ treatment. Scale bars = 50 μm. Images in the lower panel are merged with bright-field images of root tips. **d**, Quantitative data show the accumulation of H_2_O_2_ in Arabidopsis root tips. The box plot shows the mean with SD. Col-0: n = 10 (mock) and n = 12 (100 μM H_2_O_2_); *pdlp5*: n =11 (mock) and n = 13 (100 μM H_2_O_2_); and *PDLP5-HF*: n = 11 (mock) and n = 14 (100 μM H_2_O_2_). Asterisks indicate statistically significant differences compared with wild-type Col-0 (*t*-Test; two-tailed; P < 0.05).

PDLP6 also significantly enriches PIPs compared to wild-type Col-0 in proximity labeling assays; however, PDLP5 enriches PIPs to a greater extent than PDLP6^23^. We therefore tested whether PDLP6 influences H_2_O_2_ transport. In contrast to PDLP5, PDLP6 does not appear to regulate H_2_O_2_ transport in root tissues: both *pdlp6* mutants and *PDLP6-HF* overexpressors displayed DCF fluorescence levels similar to Col-0 following exogenous H_2_O_2_ treatment (Extended Data Fig. 3).

### PDLP5 represses root growth inhibition by H_2_O_2_ in Arabidopsis

Exogenous application of H_2_O_2_ is known to inhibit root growth in Arabidopsis^43–45^. Given that PDLP5 negatively regulates H_2_O_2_ uptake, we examined whether it influences root growth inhibition in response to H_2_O_2_ treatment. As expected, *PDLP5-HF* seedlings exhibited enhanced tolerance to H_2_O_2_, showing significantly longer primary roots than Col-0 under H_2_O_2_ treatment (Fig. 4). In contrast, *pdlp5* mutants were hypersensitive, exhibiting shorter primary roots compared to Col-0 (Fig. 4). These findings indicate that PDLP5 modulates H_2_O_2_-induced root growth inhibition, likely by restricting H_2_O_2_ entry into cells. Consistent with results from the H_2_O_2_ transport assay (Extended Data Fig. 3), PDLP6 did not influence root growth responses to H_2_O_2_, as Col-0, *pdlp6* knockout mutants, and *PDLP6-HF* overexpressor displayed comparable root lengths following treatment (Extended Data Fig. 4).

**Fig. 4:**
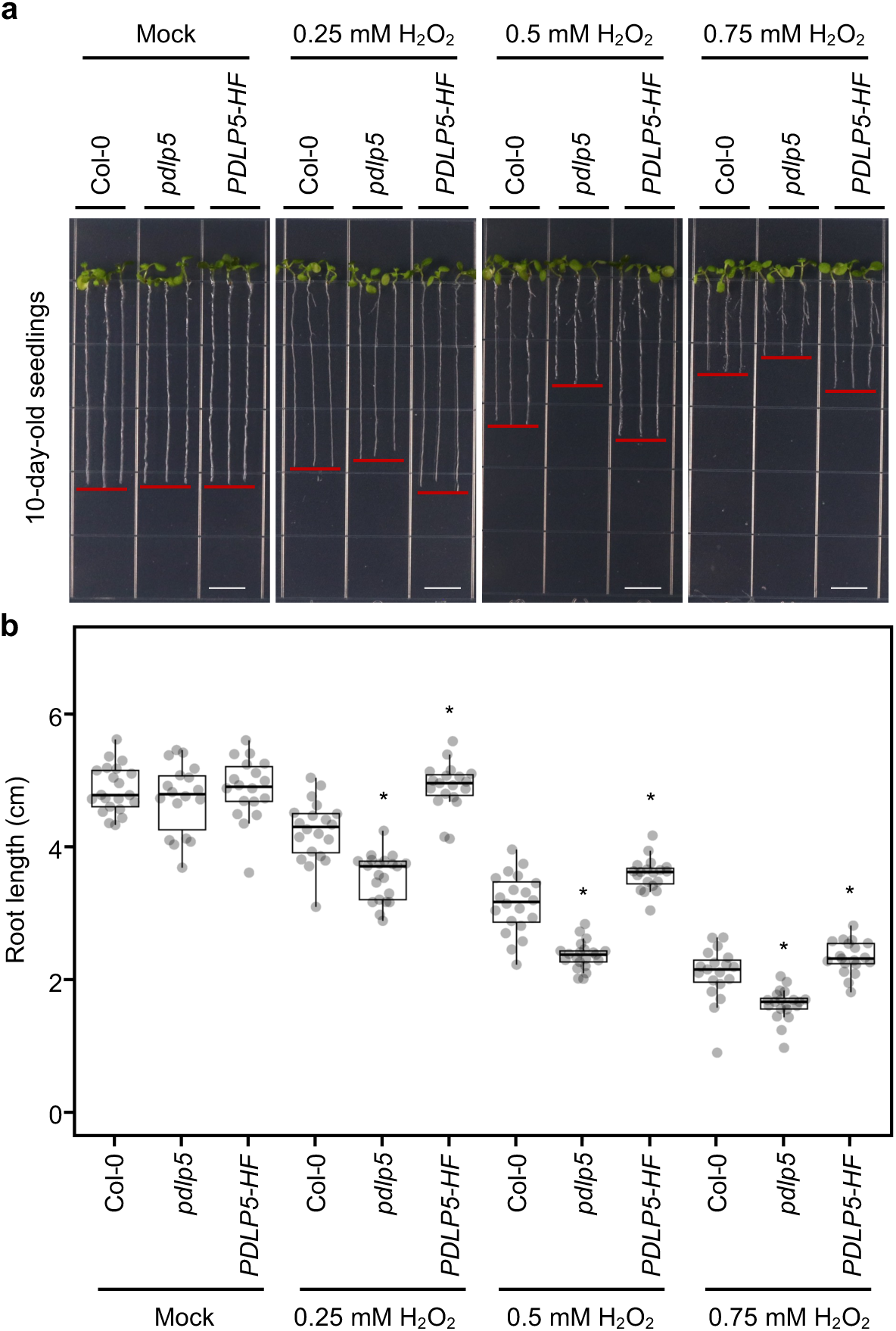
PDLP5 modulates H_2_O_2_-induced root growth inhibition in Arabidopsis. **a,** Pictures show the inhibition of root growth by H_2_O_2_. Arabidopsis seeds were sown on ½ LS agar plates containing 0.25 mM, 0.5 mM, or 0.75 mM H_2_O_2_. Root lengths were measured 10 days after germination. Scale bars = 1 cm. **b,** Quantitative data show root inhibition by H_2_O_2_. The box plot shows the mean with SD. Col-0: n = 21 (mock), n = 20 (0.25 mM H_2_O_2_), n = 20 (0.5 mM H_2_O_2_), and n = 19 (0.75 mM H_2_O_2_); *pdlp5*: n = 18 (mock), n=21 (0.25 mM H_2_O_2_), n = 21 (0.5 mM H_2_O_2_), and n = 20 (0.75 mM H_2_O_2_); and *PDLP5-HF*: n = 18 (mock), n = 19 (0.25 mM H_2_O_2_), n = 19 (0.5 mM H_2_O_2_), and n = 19 (0.75 mM H_2_O_2_). Asterisks indicate statistically significant differences compared with wild-type Col-0 (*t*-Test; two-tailed; P < 0.05).

### PIP2;5 is a critical component of PDLP5-mediated regulation of H_2_O_2_ transport

PIPs are known for their PM localization and are commonly used as PM markers^46^. Transient overexpression of all 13 sfGFP-PIP in *N. benthamiana* resulted in fluorescent signals predominantly at the PM, with infrequent signals also detected at the ER and at the aniline blue-stained callose marking plasmodesmata (Extended Data Fig. 5). All Arabidopsis PIPs transport H_2_O_2_ in yeasts^39^. To identify which PIP(s) could contribute to H_2_O_2_ uptake in root tips, we screened ten available PIP mutants representing six PIPs for their responses to H_2_O_2_. Among them, two independent lines of *pip2;5* mutants, *pip2;5-2* and *pip2;5-3*, exhibited significantly reduced DCF fluorescence in root tips compared with Col-0 upon exogenous H_2_O_2_ treatment, indicating decreased H_2_O_2_ uptake (Figs. 5a and 5b; Extended Data Figs. 6a and 6b). Consistent with this, the *pip2;5* mutants showed increased tolerance to H_2_O_2_, as evidenced by significantly longer roots than Col-0 across multiple H_2_O_2_ concentrations (Figs. 5c and 5d; Extended Data Figs. 6c and 6d). In addition, *pip1;2* mutants displayed modestly enhanced tolerance at low H_2_O_2_ concentrations, suggesting a minor role for PIP1;2 in mediating H_2_O_2_ uptake in roots (Extended Data Fig. 7).

**Fig. 5:**
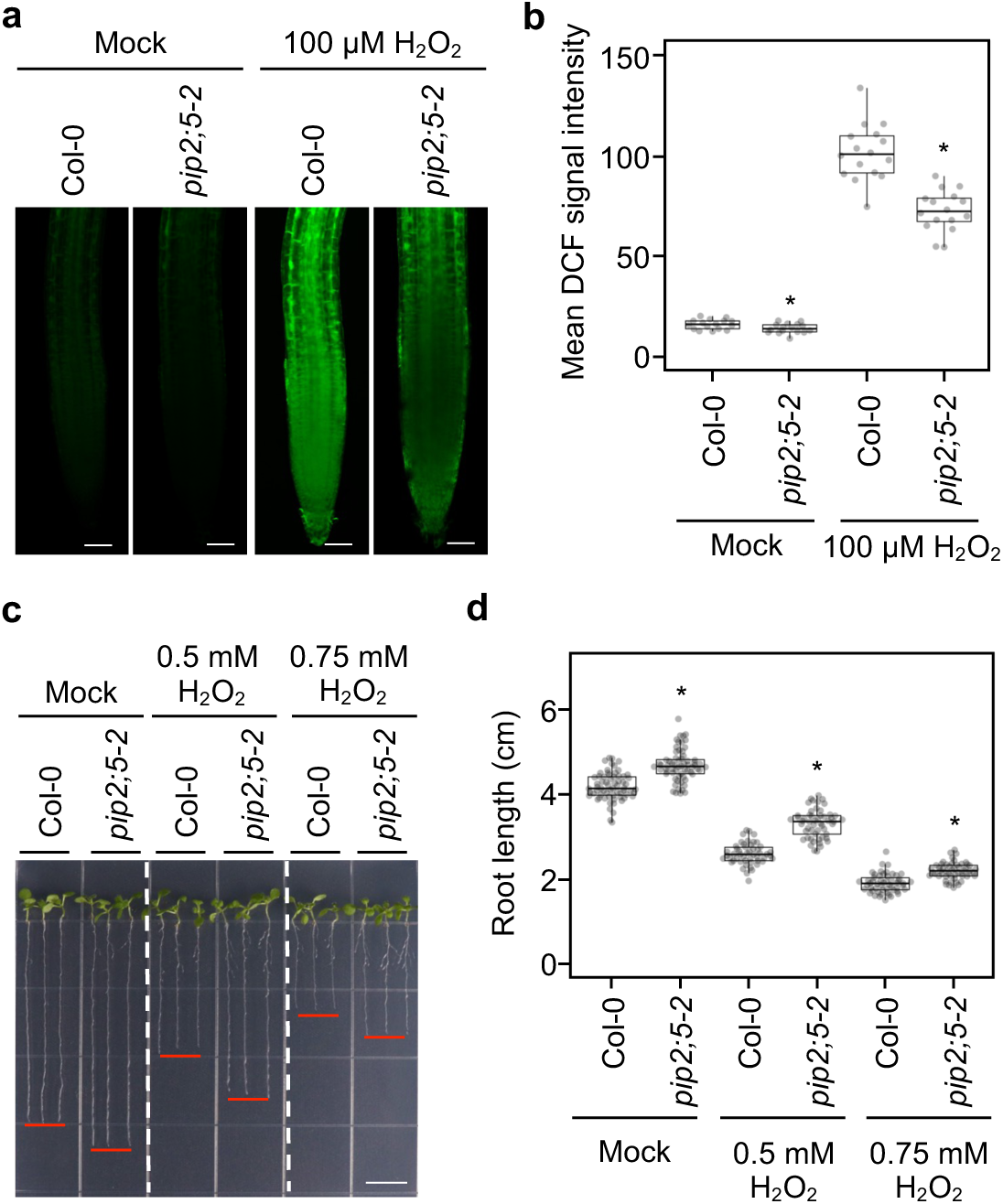
PIP2;5 plays a major role in transporting H_2_O_2_ into Arabidopsis root tips. **a,** Confocal images show the uptake of H_2_O_2_ in Arabidopsis root tips. 7-day-old Arabidopsis seedlings were incubated with 10 mM H_2_DCFDA for 15 minutes prior to H_2_O_2_ treatment. DCF signals were imaged 10 minutes after the H_2_O_2_ treatment. Scale bars = 50 μm. **b,** Quantitative data show the accumulation of H_2_O_2_ in Arabidopsis root tips. The box plot shows the mean with SD. n=16. Asterisks indicate statistically significant differences compared with wild-type Col-0 (*t*-Test; two-tailed; P < 0.05). **c,** Pictures show root inhibition by H_2_O_2_. Arabidopsis seeds were sown on ½ LS agar plates containing 0.5 mM or 0.75 mM H_2_O_2_. Root lengths were measured 10 days after germination. Scale bars = 1 cm. **d,** Quantitative data show root inhibition by H_2_O_2_. The box plot shows the mean with SD. Col-0: n = 66 (mock), n = 50 (0.5 mM H_2_O_2_), and n = 53 (0.75 mM H_2_O_2_); *pip2;5-2*: n = 64 (mock), n = 53 (0.5 mM H_2_O_2_), and n = 53 (0.75 mM H_2_O_2_). Asterisks indicate statistically significant differences compared with wild-type Col-0 (*t*-Test; two-tailed; P < 0.05).

**Fig. 6:**
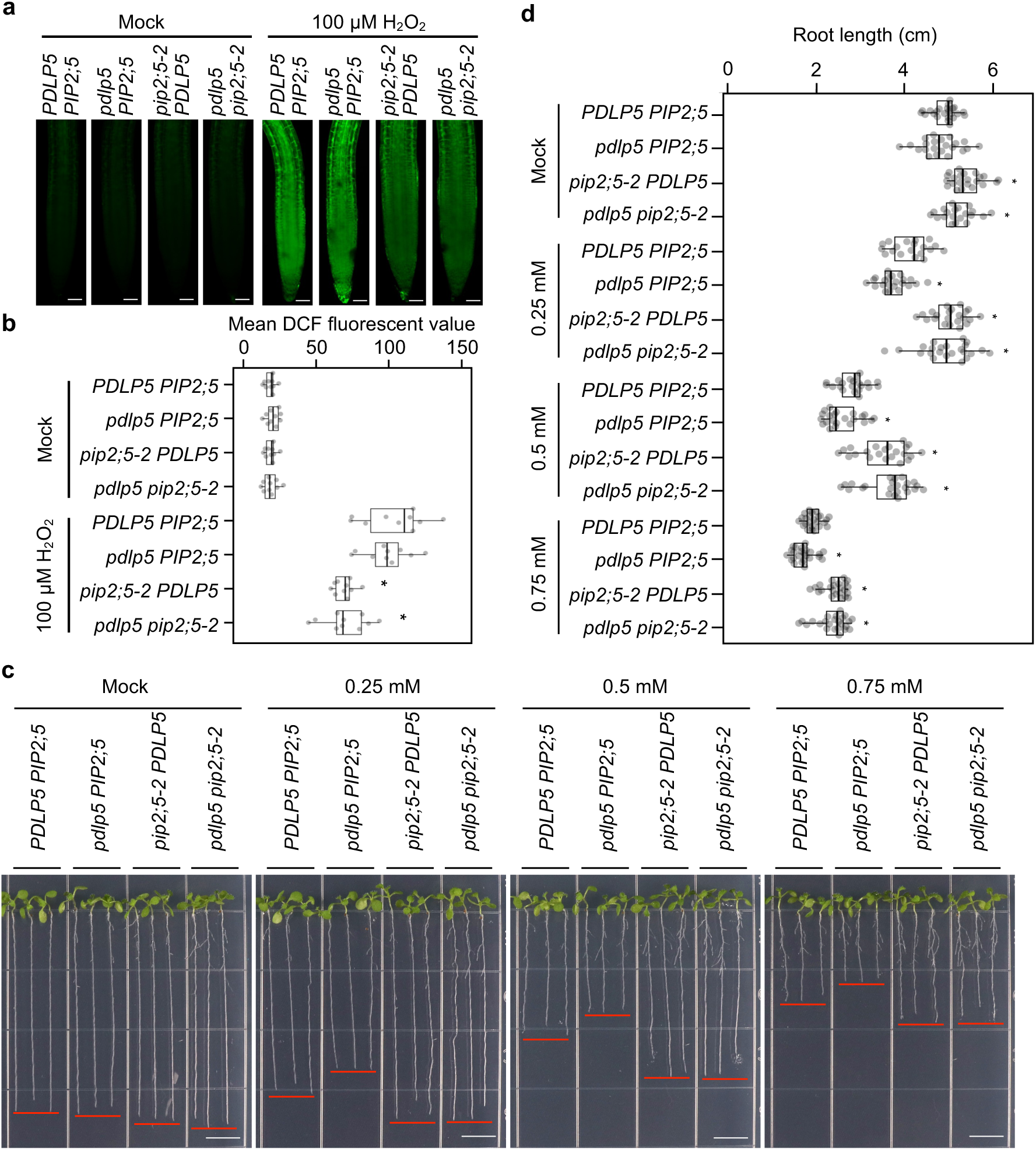
PIP2;5 is a critical component of PDLP5-mediated regulation of H_2_O_2_ transport. **a,** Confocal images show the uptake of H_2_O_2_ in Arabidopsis root tips. 7-day-old Arabidopsis seedlings were incubated with 10 mM H_2_DCFDA for 15 minutes prior to H_2_O_2_ treatment. DCF signals were imaged 10 minutes after the H_2_O_2_ treatment. Scale bars = 50 μm. **b,** Quantitative data show the accumulation of H_2_O_2_ in Arabidopsis root tips. The box plot shows the mean with SD. n = 10. Asterisks indicate statistically significant differences compared with azygous *PDLP5 PIP2;5* (*t*-Test; two-tailed; P < 0.05). **c**, Pictures show root inhibition by H_2_O_2_. Arabidopsis seeds were sown on ½ LS agar plates containing 0.25mM, 0.5 mM, or 0.75 mM H_2_O_2_. Root lengths were measured 10 days after germination. Scale bars = 1 cm. **d,** Quantitative data show root inhibition by H_2_O_2_. The box plot shows the mean with SD. *PDLP5 PIP2;5*: n = 23 (mock), n=19 (0.25 mM H_2_O_2_), n = 21 (0.5 mM H_2_O_2_), and n = 24 (0.75 mM H_2_O_2_); *pdlp5 PIP2;5*: n = 22 (mock), n= 22 (0.25 mM H_2_O_2_), n = 22 (0.5 mM H_2_O_2_), and n = 24 (0.75 mM H_2_O_2_); *pip2;5-2 PDLP5*: n = 22 (mock), n= 22 (0.25 mM H_2_O_2_), n = 22 (0.5 mM H_2_O_2_), and n = 23 (0.75 mM H_2_O_2_). *pdlp5 pip2;5-2*: n = 20 (mock), n= 23 (0.25 mM H_2_O_2_), n = 23 (0.5 mM H_2_O_2_), and n = 24 (0.75 mM H_2_O_2_). Asterisks indicate statistically significant differences compared with azygous *PDLP5 PIP2;5* (*t*-Test; two-tailed; P < 0.05).

To determine whether PIP2;5 is required for PDLP5-mediated regulation of H_2_O_2_ transport, we generated a *pdlp5 pip2;5* double mutant by crossing *pdlp5* with *pip2;5-2*. From the F_2_ progeny, we identified *PDLP5 PIP2;5* as the azygous, *pdlp5 PIP2;5* and *pip2;5-2 PDLP5* as the single mutants, and *pdlp5 pip2;5-2* as the double mutant (Extended Data Fig. 8). We then examined the H_2_O_2_ uptake in the root tip using H_2_DCFDA staining. Compared to *PDLP5 PIP2;5*, *pip2;5-2 PDLP5* and *pdlp5 pip2;5-2* double mutants displayed significantly reduced DCF fluorescence (Figs. 6a and 6b). Similarly, *pip2;5-2 PDLP5* and *pdlp5 pip2;5-2* plants exhibited longer roots than *PDLP5 PIP2;5* and *pdlp5 PIP2;5* (Figs. 6c and 6d). Consistent with previous observations (Fig. 4), *pdlp5 PIP2;5* mutants are more sensitive to root growth inhibition by H_2_O_2_ (Figs. 6c and 6d). Together, the *pdlp5 pip2;5-2* double mutant phenocopied *pip2;5-2 PDLP5*, indicating that PIP2;5 is required for PDLP5 to regulate H_2_O_2_ transport.

## Discussion

Using complementary approaches, we show that PDLP5 associates with PIPs and restricts H_2_O_2_ uptake in a PIP2;5-dependent manner. Given the central role of AQPs in facilitating the transport of water, H_2_O_2_, and other small neutral molecules, their activity is tightly regulated. In plants, PIP function is primarily modulated by phosphorylation of cytosolic termini and intracellular loops^10,47-^_53_, as well as by interactions with cytosolic proteins, as observed for animal AQPs^54^. Our findings reveal a distinct mechanism in which PDLP5, through its ectodomain, interacts with the extracellular face of PIP channels, acting as a physical modulator that suppresses channel activity from the apoplastic side.

AlphaFold3 predicted not only PDLP5-PIP heterodimers but also higher-order PDLP5-PIP and PDLP5-HsAQP assemblies (Extended Data Fig. 1), in which the PDLP5 ectodomain is positioned to engage the extracellular surface of AQP channels from different organisms. Together with in vivo and genetic data, these models provide a compelling framework for understanding how PDLP5 regulates the function of AQPs. The physical association between PDLP5 and PIPs, combined with PDLP5-dependent restriction of H_2_O_2_ uptake and enhanced oxidative tolerance in PDLP5 overexpressor and *pip2;5* mutants, supports a model in which PDLP5 acts as an extracellular gatekeeper that limits PIP-mediated H_2_O_2_ diffusion across the PM. Resolving the PDLP5-PIP interface at high resolution will be essential for validating this mechanism, a previously unrecognized mode of extracellular control over redox signaling and membrane transport in plants.

PDLP5 is a key regulator of plasmodesmal function, promoting the accumulation of callose at plasmodesmata^20^. Beyond this local role, PDLP5 contributes to systemic calcium and ROS signaling during stress responses^38,55^ and has emerged as a central modulator of plant growth, development, and immunity^20,21–24,27^. During pathogen perception, plant cells produce eROS, including H_2_O_2_, through PM-localized RBOHs^56,57^. This eROS burst both restricts pathogen spread and reinforces the cell wall through oxidative cross-linking, while also serving as a signal perceived by sensors such as CARD1/HPCA1^44,58,59^. Controlled H_2_O_2_ transport from the apoplast to the cytosol via PIPs is therefore essential for proper intracellular ROS signaling^9,10,38,40^. We propose that PDLP5 modulates this process by associating with PIPs to maintain redox homeostasis between the apoplast and cytosol, restricting excessive H_2_O_2_ influx while sustaining apoplastic pools necessary for local defense. Such regulation would spatially and temporally confine ROS signaling, limiting oxidative damage while ensuring robust immune activation.

Although PIPs are typically considered components of the bulk PM, several members have been detected in plasmodesmal proteomes^60^ and identified as interacting proteins of plasmodesmata-associated proteins^22,23,61,62^ (Fig. 1). Consistent with these findings, transient expression in *N. benthamiana* revealed partial plasmodesmal localization of PIPs (Extended Data Fig. 5), suggesting that a subset may also operate at the PD-PM. While PDLP5 is largely plasmodesmata-specific, it may also act at the bulk PM, as PDLP5-YFP signals have been observed at the PM both when expressed under a constitutive promoter^22,23,63^ and under its native promoter^23^. Resolving the spatial context in which PDLP5-PIP complexes operate will clarify the link between aquaporin-mediated ROS transport and the regulation of intercellular communication.

When expressed in yeast, PIP1 isoforms from *Arabidopsis* and maize predominantly accumulate in the ER, whereas PIP2;5 proteins are efficiently targeted to the PM. Co-expression of PIP2;5 with its cognate PIP1 facilitates the localization of PIP1 to the PM, indicating that PIP2;5 assists in PIP1 trafficking^39,64–66^. These findings raise the possibility that PIP2;5 plays a similar role in *Arabidopsis*, supporting the proper PM and potentially plasmodesmal localization of PIP1 isoforms. Consistent with this notion, *pip2;5* mutants exhibit reduced H_2_O_2_ uptake and attenuated sensitivity to H_2_O_2_-induced root growth inhibition (Fig. 5 and Extended Data Fig. 6). Future work examining PIP1 localization in the *pip2;5* background will help clarify whether PIP2;5 serves as a chaperone or assembly factor required for PIP1 recruitment to the PM.

Our findings reveal an extracellular mechanism for regulating AQPs, establishing a direct link between plasmodesmal signaling and cellular redox homeostasis. While PDLP5 is specific to land plants^63^, its ectodomain can interact with HsAQPs (Extended data Fig. 2), suggesting the potential for PDLP5 to inhibit AQPs across kingdoms. Because AQPs also mediate water transport, PDLP5 may influence host-pathogen interactions by modulating water flux, a potential strategy exploited by microbes during infection. More broadly, this study provides a conceptual framework for extracellular regulation of AQPs, with implications for enhancing crop resilience and informing therapeutic strategies targeting AQPs in diverse biological systems.

## Methods

### Plant Material and Growth Conditions

*Arabidopsis thaliana* (*Arabidopsis*) and *Nicotiana benthamiana* plants were grown at 22 °C with 50% humidity and irradiated with 110 µmol m^-2^ s^-1^ of white light for 16 hours per day. Arabidopsis wild type (Col-0), *pdlp5, PDLP5-HF, pdlp6-1, pdlp6-2,* and *PDLP6-HF* were reported previously (Li et al., 2024). *pip1;2-1* (SALK_019794C), *pip1;2-2* (SALK_145347C), *pip2;5-2* (SAIL_452_H09), and *pip2;5-3* (SM_3_30592) were obtained from ABRC (Columbus, OH). Primers used for genotyping are listed in Table S1. For seedlings grown on plates, seeds were sterilized in 75% ethanol and germinated on ½ × Linsmaier & Skoog (LS) medium plates containing 1% sucrose and 1% Bacto agar.

### Gene Cloning and Plasmid Construction

Constructs were generated using the standard Gateway cloning system (Invitrogen). Primers and gBlock DNA fragments used for cloning are listed in Table S1, and vectors used in this study are listed in Table S2. Coding sequences of *PIPs* were amplified with Gateway-compatible primers from cDNA synthesized from total RNA extracted from wild-type Col-0 seedlings. Coding sequences of *HsAQP*s were synthesized as gBlock Gene Fragments (IDT), except for *HsAQP9* (Addgene plasmid #48808), which was amplified from plasmid DNAs order from Addgene. For split-ubiquitin yeast two-hybrid (suY2H) assays (Bashline et al., 2015), mbSUS Gateway destination vector *MetYC_GW* was used to fuse CubPLV peptide to the C-terminus of PDLP5 (PDLP5-Cub), and *NX32_GW* was used to fuse PIPs and HsAQPs with the NubG peptide at the N-terminus (NubG-PIPs and NubG-HsAQPs). For co-immunoprecipitation (Co-IP) assays, sfGFP-tagged PIPs were cloned into *pEG100-35S-sfGFP-GW*. For luciferase complementation assays, *pGWB-GW-nLUC* (Addgene Plasmid #174050) was used to generate PDLP5-nLuc, and *pGWB-cLUC-GW* (Addgene Plasmid #174051) was used for cLuc-PIPs, cLuc-HsAQPs, cLuc-TET1, and cLuc-YFP.

### AlphaFold3 structural predictions

PDLP5 heterodimer and octamer complexes with AtPIPs and HsAQPs were predicted using AlphaFold3 (https://alphafoldserver.com/). The signal peptide of PDLP5 was truncated prior to prediction.

### Split-ubiquitin yeast two-hybrid (suY2H) assays

suY2H assays were performed as previously described^67^ with minor modifications. The yeast host strain THY.AP4 (*MATa ura3 leu2 lexA::lacZ::trp1 lexA::HIS3 lexA::ADE2*) was used for transformation with the Frozen-EZ Yeast Transformation II Kit (Zymo Research, #T2001). Fresh overnight cultures of THY.AP4 grown in YPAD medium at 30 °C were adjusted to an OD600 of 0.8-1.0 to prepare competent cells. Cells were pelleted at 500 × g for 4 min, washed with 10 mL of EZ1 solution, and resuspended in 1 mL of EZ2 solution. Aliquots of 50 µL were frozen at −80 °C in a styrofoam container for gradual cooling. For transformation, 6 µL of competent cells were mixed with 0.12-0.6 µg of plasmid DNA, followed by the addition of 60 µl of EZ3 solution. After mixing, the cells were incubated at 30 °C for 60-90 min, with vigorous mixing performed 2-3 times during the incubation period. Transformants were spread on selective plates (SD−Leu−Trp) and incubated at 30 °C for 2-4 days. Six colonies were picked and streaked onto selective plates, as well as selective plates supplemented with X-gal, and incubated for an additional 2-4 days until blue coloration developed. One representative clone per transformant was selected for liquid culture in selective medium at 30 °C overnight. Cultures were adjusted to OD600 values of 1 and 0.1, and 5 µl of each dilution was spotted onto three selective media: (i) SD−Leu−Trp (−LW) + X-gal, (ii) SD−Leu−Trp−His (−LWH) + Met, and (iii) SD−Leu−Trp−His (−LWH) + 3-AT. Plates were incubated at 30 °C for 2-4 days before imaging.

### Co-Immunoprecipitation (CoIP) and Immunoblot Analyses

*Agrobacterium tumefaciens* GV3101 strains carrying the plasmid(s) of interest were adjusted to an optical density at 600 nm (OD_600_) of 0.2 and infiltrated into fully expanded *N. benthamiana* leaves. Infiltrated tissues were collected 2 d post-infiltration. CoIP assays were performed as described previously^27^. Briefly, infiltrated tissues were harvested, flash-frozen, and ground in liquid nitrogen, then lysed in 3 ml of RIPA buffer (50 mM Tris-HCl, pH 7.5, 150 mM NaCl, 1% NP-40, 1% sodium deoxycholate, 0.1% SDS, and 1× protease inhibitor cocktail, Roche). Lysates were rotated for 1 h at 4 °C and centrifuged at 13,000 × g for 10 min. The supernatants were filtered through two layers of Miracloth and then centrifuged again. A 60-µl aliquot was mixed with 20 µl of 4× SDS loading buffer (Bio-Rad) and heated at 70 °C for 10 min to generate the input sample. The remaining supernatant was incubated overnight at 4 °C with 15 µl of pre-washed GFP-Trap® Magnetic Agarose (ChromoTek). Beads were collected on a magnetic rack, washed four times with RIPA buffer, and resuspended in 50 µL of 2× SDS sample buffer. They were then heated at 70 °C for 10 min. After centrifugation, proteins in the supernatant were analyzed by immunoblotting.

For immunoblot analyses, proteins were separated on 4-15% Mini-PROTEAN TGX precast gels (Bio-Rad) and transferred to polyvinylidene fluoride membranes (Bio-Rad) using the Trans-Blot Turbo Transfer System with the RTA transfer kit (Bio-Rad). Membranes were blocked in 3% (w/v) BSA in TBST (50 mM Tris base, 150 mM NaCl, 0.05% [v/v] Tween 20, pH 8.0) for 1 h at room temperature, then incubated overnight at 4 °C with either α-GFP (1:10,000; Abcam, ab290) or α-Flag-HRP (1:10,000; Sigma-Aldrich, A8592) diluted in blocking buffer. Membranes were washed four times in TBST for 10 min each. For GFP detection, membranes were incubated with goat anti-rabbit IgG secondary antibody (1:20,000; Thermo Fisher Scientific, 31460), followed by four additional washes in TBST. Signals were visualized using Clarity or Clarity Max Western ECL.

### Luciferase complementation assay

Luciferase complementation assays were performed as described previously^68^ with minor modifications. *Agrobacterium tumefaciens* GV3101, carrying the constructs of interest, was grown overnight at 30 °C in LB medium supplemented with the appropriate antibiotics. Cells from 1 mL of culture were collected by centrifugation at 4,000 rpm for 5 min at room temperature and then resuspended in 1 mL of induction solution containing 150 µM acetosyringone. After a second centrifugation step, pellets were resuspended in induction solution and adjusted to an OD_600_ of 0.5. Cultures were incubated at 30 °C for 2 h without shaking, mixed in equal volumes of different bacterial strains, and infiltrated into fully expanded *N. benthamiana* leaves using a needleless 1 ml syringe. Plants were maintained under growth chamber conditions for 48 h prior to luminescence measurements. For imaging, infiltrated leaves were excised, placed on trays, and sprayed with 1 mM D-luciferin (Invitrogen, L2916) until fully wetted. Samples were incubated in darkness for 10 min, and luminescence was captured using an iBright imaging system (Chemic Blot mode). For quantitative measurements, 0.4-cm-diameter leaf discs were collected from infiltrated regions using a biopsy punch, placed in a 96-well plate containing 1 mM D-luciferin, incubated for 10 min in darkness, and measured using a plate reader (BioTek, Synergy H1).

### Subcellular localization assay

*Agrobacterium tumefaciens* GV3101 strains carrying the plasmid of interest were adjusted to an OD_600_ of 0.2 and infiltrated into fully expanded *N. benthamiana* leaves^69^. Infiltrated tissues were imaged by confocal microscopy 2 d post-infiltration.

### Protoplast H_2_O_2_ uptake assay

Arabidopsis leaf protoplasts were isolated from 4-week-old plants as previously described^70^. Protoplasts were incubated overnight in WI solution containing 500 mM D-mannitol. Protoplasts were transferred into a 12-well plate, and 10 μM H_2_DCFDA and 100 μM H_2_O_2_ were added to each well. Plates were incubated for 20 min. DCF fluorescence was imaged by confocal microscopy, and signal intensities were quantified using Fiji.

### Root H_2_O_2_ uptake assay

Sterilized Arabidopsis seeds were sown on ½ LS medium plates, stratified at 4 °C for 2 days, and then transferred to a growth chamber for vertical growth. Seven-day-old seedlings were transferred into 6-well plates containing 4 mL of ½ LS medium solution with 1% sucrose per well. Seedlings were incubated with 10 µM H_2_DCFDA for 15 min, followed by treatment with 100 µM H_2_O_2_ for 10 min. The seedlings were then washed with a ½ LS medium solution containing 1% sucrose. DCF fluorescence in root tips was imaged using confocal microscopy, and fluorescence intensities were quantified using Fiji.

### Root H_2_O_2_ sensitivity assay

Sterilized seeds were sown on ½ LS medium plates supplemented with 0 mM, 0.25 mM, 0.5 mM, and 0.75 mM H_2_O_2_, stratified at 4 °C for 2 days and then transferred to a growth chamber for vertical growth. After 10 days, the seedlings were images, and the root length was measured by Fiji.

### ROS burst assay

ROS burst assays were performed using leaf discs from fully expanded mature Arabidopsis leaves. Leaf discs (4 mm diameter) were collected with a biopsy punch from the middle region of the leaf, avoiding the mid-vein, and incubated adaxial side up in 150 μl ddH_2_O in a 96-well luminometer plate for 12 h at room temperature. The water was carefully removed and replaced with 100 μl of reaction solution containing 20 μg/mL horseradish peroxidase (HRP), 50 μM luminol L-012, and 100 nM flg22 (mock treatment with ddH_2_O). Luminescence was measured using a microplate reader (BioTek, Synergy H1) with kinetic measurements for 60 minutes, with readings taken every 2 min. Relative light units were recorded, and data were processed to generate time-course plots of ROS production.

### AUR staining assay

The 4^th^ leaves from 3-week-old plants were infiltrated with 100 nM flg22 (mock treatment with ddH_2_O) and 10 μM AUR. Samples were mounted in AUR solution on slides 30 min later and the fluorescent signal was detected using confocal microscopy. The relative signal intensity was measured by Fiji.

### Confocal Imaging

All confocal images were acquired using a Zeiss LSM 700 laser-scanning microscope. YFP and sfGFP were excited at 488 nm, and the emission was collected using a SP555 filter. DCF was excited at 488 nm, with emission collected using a SP555 filter. The fluorescent AUR product was excited at 555 nm, and emission was collected using a SP640 filter. Aniline blue-stained callose was excited at 405 nm, and emission was collected using a SP490 filter. FM4-64 was excited at 405 nm, and emission was collected using an LP 640 filter.

## Statistical Analysis

Column plots were created using Excel. Box plots were created with an online software (https://huygens.science.uva.nl/PlotsOfData/). Student’s *t*-test was performed to test the statistical significance of differences.

## Data availability

Data supporting the findings of this study are included in the main article and its Supplementary Information. Source data are provided with this paper. All other datasets generated or analyzed during the study are available from the corresponding author upon request.

## Acknowledgements

We thank Dr. Dipali Sashital (Iowa State University) for introducing us to AlphaFold and for critically reviewing the manuscript. This work was supported by the National Science Foundation (grant MCB2339067 to K.A.), the ISU Crop Bioengineering Center (K.A.), and the ISU Frontier Science Fund (K.A.).

## Author Information

Authors and Affiliations

**Department of Genetics, Development, and Cell Biology, Iowa State University, Ames, Iowa, USA**

Zhongpeng Li, Su-Ling Liu, Saiful Islam, Madelyn Clements, Yani Chen & Kyaw Aung

## Contributions

Z.L., S.L., S.I., and K.A. designed the experiments. Z.L. performed the majority of the experiments and carried out data analysis. S.L. designed and generated vectors and plasmids and conducted the suY2H assays. S.I. performed the AlphaFold analyses and assisted with independent confirmation of some results. M.C. and Y.C. contributed to screening the *pip* mutants. Z.L. and K.A. wrote the manuscript with input from all authors. K.A. supervised the study.

Corresponding authors

Correspondence to Kyaw Aung.

## Ethic declarations

### Competing interests

The authors declare no competing interests.

**Extended Data Fig. 1:** AlphaFold predicts PDLP5-AQP complexes. **a**, AlphaFold3 predicts the formation of a PDLP5-PIP1;1 heterodimer; the PDLP5 signal peptide was omitted from the prediction. **b**, AlphaFold3 predicts the formation of a PDLP5-PIP1;1 hetero-octamer. **c**, AlphaFold3 predicts a PDLP5-HsAQP0 heterodimer. Predicted local distance difference test (pLDDT) scores range from 0 (lowest confidence) to 100 (highest confidence), whereas predicted template modeling (pTM) and interfacial predicted template modeling (ipTM) scores range from 0 (lowest confidence) to 1 (highest confidence).

**Extended Data Fig. 2:** PDLP5 physically interacts with human aquaporins (HsAQPs). **a**, Split-ubiquitin yeast two-hybrid assay determines the interaction between PDLP5 and HsAQPs. Nub-WT and NubG were used as positive and negative controls, respectively. To conduct blue/white screening, -LW + X-Gal synthetic dropout plates were utilized, while -LWH synthetic dropout plates were used for histidine auxotrophy selection. To increase the stringency of the assay, 30 mM 3-AT or 25 mM Met was added to the -LWH synthetic dropout plates. **b**, The luciferase complementation assay demonstrates the interaction between PDLP5 and HsAQPs. Agrobacteria carrying the respective plasmids were co-infiltrated into *N. benthamiana* leaves. Two days after infiltration, leaves were sprayed with 1 mM D-luciferin, and luminescence was imaged using an iBright system (top). **c**, Quantification of luminescence was performed using a microplate reader (bottom). Data represent the mean ± SD (n = 5).

**Extended Data Fig. 3:** PDLP6 does not affect H_2_O_2_ transport in Arabidopsis root tips. **a**, Confocal images show the uptake of H_2_O_2_ in Arabidopsis root tips. 7-day-old Arabidopsis seedlings were incubated with 10 mM H_2_DCFDA for 15 minutes prior to H_2_O_2_ treatment. DCF signals were imaged 10 minutes after the H_2_O_2_ treatment. Images in the lower panel are merged with bright-field images of root tips. Scale bars = 50 μm. **b**, Quantitative data show the accumulation of H_2_O_2_ in Arabidopsis root tips. The box plot shows the mean with SD. Col-0: n = 17 (mock) and n = 15 (100 μM H_2_O_2_); *pdlp6-1*: n = 17 (mock) and n = 20 (100 μM H_2_O_2_); *pdlp6-2*: n = 17 (mock) and n = 17 (100 μM H_2_O_2_); and *PDLP6-HF*: n = 14 (mock) and n = 19 (100 μM H_2_O_2_).

**Extended Data Fig. 4:** PDLP6 does not modulate H_2_O_2_-induced inhibition of root growth in Arabidopsis. **a**, Pictures show root inhibition by H_2_O_2_. Arabidopsis seeds were sown on ½ LS agar plates containing 0.25 mM, 0.5 mM or 0.75 mM H_2_O_2_. Root lengths were measured 10 days after germination. Scale bar = 1 cm. **b**, Quantitative data show root inhibition by H_2_O_2_. The box plot shows the mean with SD. Col-0: n = 29 (mock), n = 32 (0.25 mM H_2_O_2_), n =29 (0.5 mM H_2_O_2_), and n = 32 (0.75 mM H_2_O_2_); *pdlp6-1*: n = 29 (mock), n = 28 (0.25 mM H_2_O_2_), n = 30 (0.5 mM H_2_O_2_), and n = 28 (0.75 mM H_2_O_2_); *pdlp6-2*: n = 29 (mock), n = 29 (0.25 mM H_2_O_2_), n = 29 (0.5 mM H_2_O_2_), and n = 28 (0.75 mM H_2_O_2_); and *PDLP6-HF*: n = 28 (mock), n = 32 (0.25 mM H_2_O_2_), n = 27 (0.5 mM H_2_O_2_), and n = 29 (0.75 mM H_2_O_2_). Asterisks indicate statistically significant differences compared with wild-type Col-0 (*t*-Test; two-tailed; P < 0.05).

**Extended Data Fig. 5:** Subcellular localization of PIPs. Agrobacteria carrying sfGFP-PIPs were infiltrated into *N. benthamiana* to transiently express the fusion proteins. The subcellular localization of sfGFP fusion protein was detected using confocal microscopy. Aniline blue stains callose at plasmodesmata. A free sfGFP was used as a control to present the expression of the protein in the cytosol and nucleus. Scale bars = 5 μm.

**Extended Data Fig. 6:** PIP2;5 plays a major role in transporting H_2_O_2_ into Arabidopsis root tips. **a**, Confocal images show the uptake of H_2_O_2_ in Arabidopsis root tips. 7-day-old Arabidopsis seedlings were incubated with 10 mM H_2_DCFDA for 15 minutes prior to H_2_O_2_ treatment. DCF signals were imaged 10 minutes after the H_2_O_2_ treatment. Scale bars = 50 μm. **b**, Quantitative data show the accumulation of H_2_O_2_ in Arabidopsis root tips. The box plot shows the mean with SD. n=16. Asterisks indicate statistically significant differences compared with wild-type Col-0 (*t*-Test; two-tailed; P < 0.05). **c**, Pictures show root inhibition by H_2_O_2_. Arabidopsis seeds were sown on ½ LS agar plates containing 0.5 mM or 0.75 mM H_2_O_2_. Root lengths were measured 10 days after germination. Scale bars = 1 cm. **d**, Quantitative data show root inhibition by H_2_O_2_. The box plot shows the mean with SD. Col-0: n = 11 (mock), n = 20 (0.5 mM H_2_O_2_), and n = 19 (0.75 mM H_2_O_2_); *pip2;5-3*: n = 12 (mock), n = 18 (0.5 mM H_2_O_2_), and n = 20 (0.75 mM H_2_O_2_). Asterisks indicate statistically significant differences compared with wild-type Col-0 (*t*-Test; two-tailed; P < 0.05).

**Extended Data Fig. 7:** PIP1;2 plays a minor role in transporting H_2_O_2_ into Arabidopsis root tips. **a**, Pictures show root inhibition by H_2_O_2_. Arabidopsis seeds were sown on ½ LS agar plates containing 0.25 mM, 0.5 mM or 0.75 mM H_2_O_2_. Root lengths were measured 10 days after germination. Scale bars = 1 cm. **b**, Quantitative data show root inhibition by H_2_O_2_. The box plot shows the mean with SD. Col-0: n =33 (mock), n = 33 (0.25 mM H_2_O_2_), n = 31 (0.5 mM H_2_O_2_), and n = 28 (0.75 mM H_2_O_2_); *pip1;2-1*: n = 33 (mock), n = 33 (0.25 mM H_2_O_2_), n = 30 (0.5 mM H_2_O_2_), and n = 33 (0.75 mM H_2_O_2_); and *pip1;2-2*: n = 28 (mock), n = 28 (0.25 mM H_2_O_2_), n = 29 (0.5 mM H_2_O_2_), and n = 29 (0.75 mM H_2_O_2_). Asterisks indicate statistically significant differences compared with wild-type Col-0 (*t*-Test; two-tailed; P < 0.05).

**Extended Data Fig. 8:** Characterization of the *pdlp5 pip2;5-2* double mutant. A genetic cross was performed between *pdlp5* and *pip2;5-2* mutants. The F_2_ population was used to isolate the following genotypes using a PCR-based genotyping approach: wild type (*PDLP5 PIP2;5*), double mutant (*pdlp5 pip2;5-2*), and single mutants (*pdlp5 PIP2;5* and *pip2;5-2 PDLP5*). **a**, Genomic DNA (gDNA) was amplified using primers flanking the T-DNA insertion site to detect wild-type alleles. T-DNA-specific primers were used to confirm the presence of the T-DNA insertion within *PDLP5* or *PIP2;5*. **b**, RT-PCR was performed to assess the transcript levels of *PDLP5* and *PIP2;5* in wild type, double mutant, and single mutants. *UBQ10* was used as an internal control. The number of PCR cycles used for amplification is indicated.

**Supplementary Table S1.** Primers and gBlock DNA fragments used in this study.

**Supplementary Table S3.** Plasmids used in this study.

## References

1. Martin RE, Postiglione AE, Muday GK. Reactive oxygen species function as signaling molecules in controlling plant development and hormonal responses. Curr Opin Plant Biol. 2022;69:102293. doi:10.1016/j.pbi.2022.102293

2. Mittler R, Zandalinas SI, Fichman Y, Van Breusegem F. Reactive oxygen species signalling in plant stress responses. Nat Rev Mol Cell Biol. 2022;23(10):663–679. doi:10.1038/s41580-022-00499-2

3. Smirnoff N, Arnaud D. Hydrogen peroxide metabolism and functions in plants. New Phytol. 2019;221(3):1197–1214. doi:10.1111/nph.15488

4. Wang P, Liu WC, Han C, Wang S, Bai MY, Song CP. Reactive oxygen species: Multidimensional regulators of plant adaptation to abiotic stress and development. J Integr Plant Biol. 2024;66(3):330–367. doi:10.1111/jipb.13601

5. Bienert GP, Chaumont F. Aquaporin-facilitated transmembrane diffusion of hydrogen peroxide. Biochim Biophys Acta. 2014;1840(5):1596–1604. doi:10.1016/j.bbagen.2013.09.017

6. Chaumont F, Tyerman SD. Aquaporins: highly regulated channels controlling plant water relations. Plant Physiol. 2014;164(4):1600–1618. doi:10.1104/pp.113.233791

7. Johanson U, Karlsson M, Johansson I, et al. The complete set of genes encoding major intrinsic proteins in Arabidopsis provides a framework for a new nomenclature for major intrinsic proteins in plants. Plant Physiol. 2001;126(4):1358–1369. doi:10.1104/pp.126.4.1358

8. Maurel C, Boursiac Y, Luu DT, Santoni V, Shahzad Z, Verdoucq L. Aquaporins in Plants. Physiol Rev. 2015;95(4):1321–1358. doi:10.1152/physrev.00008.2015

9. Tian S, Wang X, Li P, et al. Plant Aquaporin AtPIP1;4 Links Apoplastic H2O2 Induction to Disease Immunity Pathways. Plant Physiol. 2016;171(3):1635–1650. doi:10.1104/pp.15.01237

10. Rodrigues O, Reshetnyak G, Grondin A, et al. Aquaporins facilitate hydrogen peroxide entry into guard cells to mediate ABA- and pathogen-triggered stomatal closure. Proc Natl Acad Sci U S A. 2017;114(34):9200–9205. doi:10.1073/pnas.1704754114

11. Zhang M, Shi H, Li N, et al. Aquaporin OsPIP2;2 links the H2O2 signal and a membrane-anchored transcription factor to promote plant defense. Plant Physiol. 2022;188(4):2325–2341. doi:10.1093/plphys/kiab604

12. Jia Z, Yu W, Guo X, et al. CPK12 decodes effector-triggered calcium signaling and phosphorylates PIP2;1 to facilitate apoplastic ROS transport into the cytoplasm in Arabidopsis. Mol Plant. 2025;18(10):1724–1741. doi:10.1016/j.molp.2025.08.018

13. Castro B, Citterico M, Kimura S, Stevens DM, Wrzaczek M, Coaker G. Stress-induced reactive oxygen species compartmentalization, perception and signalling. Nat Plants. 2021;7(4):403–412. doi:10.1038/s41477-021-00887-0

14. Lee J, Han M, Shin Y, Lee JM, Heo G, Lee Y. How Extracellular Reactive Oxygen Species Reach Their Intracellular Targets in Plants. Mol Cells. 2023;46(6):329–336. doi:10.14348/molcells.2023.2158

15. Bayer EM, Benitez-Alfonso Y. Plasmodesmata: Channels Under Pressure. Annu Rev Plant Biol. 2024;75(1):291–317. doi:10.1146/annurev-arplant-070623-093110

16. Tee EE, Faulkner C. Plasmodesmata and intercellular molecular traffic control. New Phytol. 2024;243(1):32–47. doi:10.1111/nph.19666

17. De Storme N, Geelen D. Callose homeostasis at plasmodesmata: molecular regulators and developmental relevance. Front Plant Sci. 2014;5:138. Published 2014 Apr 21. doi:10.3389/fpls.2014.00138

18. Pérez-Sancho J, Smokvarska M, Dubois G, et al. Plasmodesmata act as unconventional membrane contact sites regulating intercellular molecular exchange in plants. Cell. 2025;188(4):958–977.e23. doi:10.1016/j.cell.2024.11.034

19. Thomas CL, Bayer EM, Ritzenthaler C, Fernandez-Calvino L, Maule AJ. Specific targeting of a plasmodesmal protein affecting cell-to-cell communication. PLoS Biol. 2008;6(1):e7. doi:10.1371/journal.pbio.0060007

20. Lee JY, Wang X, Cui W, et al. A plasmodesmata-localized protein mediates crosstalk between cell-to-cell communication and innate immunity in Arabidopsis. Plant Cell. 2011;23(9):3353–3373. doi:10.1105/tpc.111.087742

21. Sager R, Wang X, Hill K, et al. Auxin-dependent control of a plasmodesmal regulator creates a negative feedback loop modulating lateral root emergence. Nat Commun. 2020;11(1):364. Published 2020 Jan 17. doi:10.1038/s41467-019-14226-7

22. Tee EE, Johnston MG, Papp D, Faulkner C. A PDLP-NHL3 complex integrates plasmodesmal immune signaling cascades. Proc Natl Acad Sci U S A. 2023;120(17):e2216397120. doi:10.1073/pnas.2216397120

23. Li Z, Liu SL, Montes-Serey C, Walley JW, Aung K. PLASMODESMATA-LOCATED PROTEIN 6 regulates plasmodesmal function in Arabidopsis vasculature. Plant Cell. 2024;36(9):3543–3561. doi:10.1093/plcell/koae166

24. Lim GH, Shine MB, de Lorenzo L, et al. Plasmodesmata Localizing Proteins Regulate Transport and Signaling during Systemic Acquired Immunity in Plants. Cell Host Microbe. 2016;19(4):541–549. doi:10.1016/j.chom.2016.03.006

25. Lee MW, Jelenska J, Greenberg JT. Arabidopsis proteins important for modulating defense responses to Pseudomonas syringae that secrete HopW1-1. Plant J. 2008;54(3):452–465. doi:10.1111/j.1365-313X.2008.03439.x

26. Wang X, Sager R, Cui W, Zhang C, Lu H, Lee JY. Salicylic acid regulates Plasmodesmata closure during innate immune responses in Arabidopsis. Plant Cell. 2013;25(6):2315–2329. doi:10.1105/tpc.113.110676

27. Aung K, Kim P, Li Z, et al. Pathogenic Bacteria Target Plant Plasmodesmata to Colonize and Invade Surrounding Tissues. Plant Cell. 2020;32(3):595–611. doi:10.1105/tpc.19.00707

28. Konstantinova N, Mor E, Verhelst E, et al. A precise balance of TETRASPANIN1/TORNADO2 activity is required for vascular proliferation and ground tissue patterning in Arabidopsis. Physiol Plant. 2024;176(1):e14182. doi:10.1111/ppl.14182

29. Vaattovaara A, Brandt B, Rajaraman S, et al. Mechanistic insights into the evolution of DUF26-containing proteins in land plants. Commun Biol. 2019;2:56. Published 2019 Feb 8. doi:10.1038/s42003-019-0306-9

30. Törnroth-Horsefield S, Wang Y, Hedfalk K, et al. Structural mechanism of plant aquaporin gating. Nature. 2006;439(7077):688–694. doi:10.1038/nature04316

31. Ho JD, Yeh R, Sandstrom A, et al. Crystal structure of human aquaporin 4 at 1.8 A and its mechanism of conductance. Proc Natl Acad Sci U S A. 2009;106(18):7437–7442. doi:10.1073/pnas.0902725106

32. Fischer G, Kosinska-Eriksson U, Aponte-Santamaría C, et al. Crystal structure of a yeast aquaporin at 1.15 angstrom reveals a novel gating mechanism. PLoS Biol. 2009;7(6):e1000130. doi:10.1371/journal.pbio.1000130

33. Verkman AS, Mitra AK. Structure and function of aquaporin water channels. Am J Physiol Renal Physiol. 2000;278(1):F13–F28. doi:10.1152/ajprenal.2000.278.1.F13

34. Abramson J, Adler J, Dunger J, et al. Accurate structure prediction of biomolecular interactions with AlphaFold 3. Nature. 2024;630(8016):493–500. doi:10.1038/s41586-024-07487-w

35. Verkman AS, Anderson MO, Papadopoulos MC. Aquaporins: important but elusive drug targets. Nat Rev Drug Discov. 2014;13(4):259–277. doi:10.1038/nrd4226

36. Kruse E, Uehlein N, Kaldenhoff R. The aquaporins. Genome Biol. 2006;7(2):206. doi:10.1186/gb-2006-7-2-206

37. Torres MA, Dangl JL, Jones JD. Arabidopsis gp91phox homologues AtrbohD and AtrbohF are required for accumulation of reactive oxygen intermediates in the plant defense response. Proc Natl Acad Sci U S A. 2002;99(1):517–522. doi:10.1073/pnas.012452499

38. Fichman Y, Myers RJ Jr, Grant DG, Mittler R. Plasmodesmata-localized proteins and ROS orchestrate light-induced rapid systemic signaling in Arabidopsis. Sci Signal. 2021;14(671):eabf0322. Published 2021 Feb 23. doi:10.1126/scisignal.abf0322

39. Groszmann M, De Rosa A, Chen W, et al. A high-throughput yeast approach to characterize aquaporin permeabilities: Profiling the Arabidopsis PIP aquaporin sub-family. Front Plant Sci. 2023;14:1078220. Published 2023 Jan 19. doi:10.3389/fpls.2023.1078220

40. Yao X, Mu Y, Zhang L, et al. AtPIP1;4 and AtPIP2;4 Cooperatively Mediate H2O2 Transport to Regulate Plant Growth and Disease Resistance. Plants (Basel). 2024;13(7):1018. Published 2024 Apr 3. doi:10.3390/plants13071018

41. Ashtamker C, Kiss V, Sagi M, Davydov O, Fluhr R. Diverse subcellular locations of cryptogein-induced reactive oxygen species production in tobacco Bright Yellow-2 cells. Plant Physiol. 2007;143(4):1817–1826. doi:10.1104/pp.106.090902

42. Akter S, Khan MS, Smith EN, Flashman E. Measuring ROS and redox markers in plant cells. RSC Chem Biol. 2021;2(5):1384–1401. Published 2021 Jun 29. doi:10.1039/d1cb00071c

43. Zhou L, Hou H, Yang T, et al. Exogenous hydrogen peroxide inhibits primary root gravitropism by regulating auxin distribution during Arabidopsis seed germination. Plant Physiol Biochem. 2018a;128:126–133. doi:10.1016/j.plaphy.2018.05.014

44. Zhang L, Wang L, Qiang X, et al. The CANNOT RESPOND TO DMBQ 1-GLUTAREDOXIN C1 module regulates Arabidopsis root growth via quinone-induced oxidation. Plant Physiol. 2025;199(2):kiaf419. Doi:10.1093/plphys/kiaf419

45. Wang Y, Liu X, Sun X, et al. The promotive and repressive effects of exogenous H2O2 on Arabidopsis seed germination and seedling establishment depend on application dose. Physiol Plant. 2025;177(1):e70098. doi:10.1111/ppl.70098

46. Nelson BK, Cai X, Nebenführ A. A multicolored set of in vivo organelle markers for co-localization studies in Arabidopsis and other plants. Plant J. 2007;51(6):1126–1136. doi:10.1111/j.1365-313X.2007.03212.x

47. Grondin A, Rodrigues O, Verdoucq L, Merlot S, Leonhardt N, Maurel C. Aquaporins Contribute to ABA-Triggered Stomatal Closure through OST1-Mediated Phosphorylation. Plant Cell. 2015;27(7):1945–1954. doi:10.1105/tpc.15.00421

48. Prado K, Cotelle V, Li G, et al. Oscillating Aquaporin Phosphorylation and 14-3-3 Proteins Mediate the Circadian Regulation of Leaf Hydraulics. Plant Cell. 2019;31(2):417–429. doi:10.1105/tpc.18.00804

49. Ai G, Xia Q, Song T, et al. A Phytophthora sojae CRN effector mediates phosphorylation and degradation of plant aquaporin proteins to suppress host immune signaling. PLoS Pathog. 2021;17(3):e1009388. Published 2021 Mar 12. doi:10.1371/journal.ppat.1009388

50. Chen Q, Liu R, Wu Y, et al. ERAD-related E2 and E3 enzymes modulate the drought response by regulating the stability of PIP2 aquaporins. Plant Cell. 2021;33(8):2883–2898. doi:10.1093/plcell/koab141

51. Lu K, Chen X, Yao X, et al. Phosphorylation of a wheat aquaporin at two sites enhances both plant growth and defense. Mol Plant. 2022;15(11):1772–1789. doi:10.1016/j.molp.2022.10.003

52. Zhang H, Yu F, Xie P, et al. A Gγ protein regulates alkaline sensitivity in crops. Science. 2023;379(6638):eade8416. doi:10.1126/science.ade8416

53. Zhu H, Bao Y, Peng H, et al. Phosphorylation of PIP2;7 by CPK28 or Phytophthora kinase effectors dampens pattern-triggered immunity in Arabidopsis. Plant Commun. 2025;6(1):101135. doi:10.1016/j.xplc.2024.101135

54. Törnroth-Horsefield S, Chivasso C, Strandberg H, et al. Insight into the Mammalian Aquaporin Interactome. Int J Mol Sci. 2022;23(17):9615. Published 2022 Aug 25. doi:10.3390/ijms23179615

55. Toyota M, Spencer D, Sawai-Toyota S, et al. Glutamate triggers long-distance, calcium-based plant defense signaling. Science. 2018;361(6407):1112–1115. doi:10.1126/science.aat7744

56. Kadota Y, Shirasu K, Zipfel C. Regulation of the NADPH Oxidase RBOHD During Plant Immunity. Plant Cell Physiol. 2015;56(8):1472–1480. doi:10.1093/pcp/pcv063

57. Morales J, Kadota Y, Zipfel C, Molina A, Torres MA. The Arabidopsis NADPH oxidases RbohD and RbohF display differential expression patterns and contributions during plant immunity. J Exp Bot. 2016;67(6):1663–1676. doi:10.1093/jxb/erv558

58. Laohavisit A, Wakatake T, Ishihama N, et al. Quinone perception in plants via leucine-rich-repeat receptor-like kinases. Nature. 2020;587(7832):92–97. doi:10.1038/s41586-020-2655-4

59. Wu F, Chi Y, Jiang Z, et al. Hydrogen peroxide sensor HPCA1 is an LRR receptor kinase in Arabidopsis. Nature. 2020;578(7796):577–581. doi:10.1038/s41586-020-2032-3

60. Fernandez-Calvino L, Faulkner C, Walshaw J, et al. Arabidopsis plasmodesmal proteome. PLoS One. 2011;6(4):e18880. Published 2011 Apr 20. doi:10.1371/journal.pone.0018880

61. Caillaud MC, Wirthmueller L, Sklenar J, et al. The plasmodesmal protein PDLP1 localises to haustoria-associated membranes during downy mildew infection and regulates callose deposition. PLoS Pathog. 2014;10(10):e1004496. Published 2014 Nov 13. doi:10.1371/journal.ppat.1004496

62. Hunter K, Kimura S, Rokka A, et al. CRK2 Enhances Salt Tolerance by Regulating Callose Deposition in Connection with PLDα1. Plant Physiol. 2019;180(4):2004–2021. doi:10.1104/pp.19.00560

63. Wang X, Robles Luna G, Arighi CN, Lee JY. An evolutionarily conserved motif is required for Plasmodesmata-located protein 5 to regulate cell-to-cell movement. Commun Biol. 2020;3(1):291. Published 2020 Jun 5. doi:10.1038/s42003-020-1007-0

64. Bienert GP, Heinen RB, Berny MC, Chaumont F. Maize plasma membrane aquaporin ZmPIP2;5, but not ZmPIP1;2, facilitates transmembrane diffusion of hydrogen peroxide. Biochim Biophys Acta. 2014;1838(1 Pt B):216–222. doi:10.1016/j.bbamem.2013.08.011

65. Jozefkowicz C, Rosi P, Sigaut L, et al. Loop A is critical for the functional interaction of two Beta vulgaris PIP aquaporins. PLoS One. 2013;8(3):e57993. doi:10.1371/journal.pone.0057993

66. Bienert MD, Diehn TA, Richet N, Chaumont F, Bienert GP. Heterotetramerization of Plant PIP1 and PIP2 Aquaporins Is an Evolutionary Ancient Feature to Guide PIP1 Plasma Membrane Localization and Function. Front Plant Sci. 2018;9:382. Published 2018 Mar 26. doi:10.3389/fpls.2018.00382

67. Bashline L, Gu Y. Using the split-ubiquitin yeast two-hybrid system to test protein-protein interactions of transmembrane proteins. Methods Mol Biol. 2015;1242:143–158. doi:10.1007/978-1-4939-1902-4_13

68. Zhou Z, Bi G, Zhou JM. Luciferase Complementation Assay for Protein-Protein Interactions in Plants. Curr Protoc Plant Biol. 2018b;3(1):42–50. doi:10.1002/cppb.20066

69. Li Z, Variz H, Chen Y, Liu SL, Aung K. Plasmodesmata-Dependent Intercellular Movement of Bacterial Effectors. Front Plant Sci. 2021;12:640277. Published 2021 Mar 22. doi:10.3389/fpls.2021.640277

70. Yoo SD, Cho YH, Sheen J. Arabidopsis mesophyll protoplasts: a versatile cell system for transient gene expression analysis. Nat Protoc. 2007;2(7):1565–1572. doi:10.1038/nprot.2007.199

