## Extended Data Fig. 1 for "PDLP5 regulates aquaporin-mediated hydrogen peroxide transport in *Arabidopsis*"

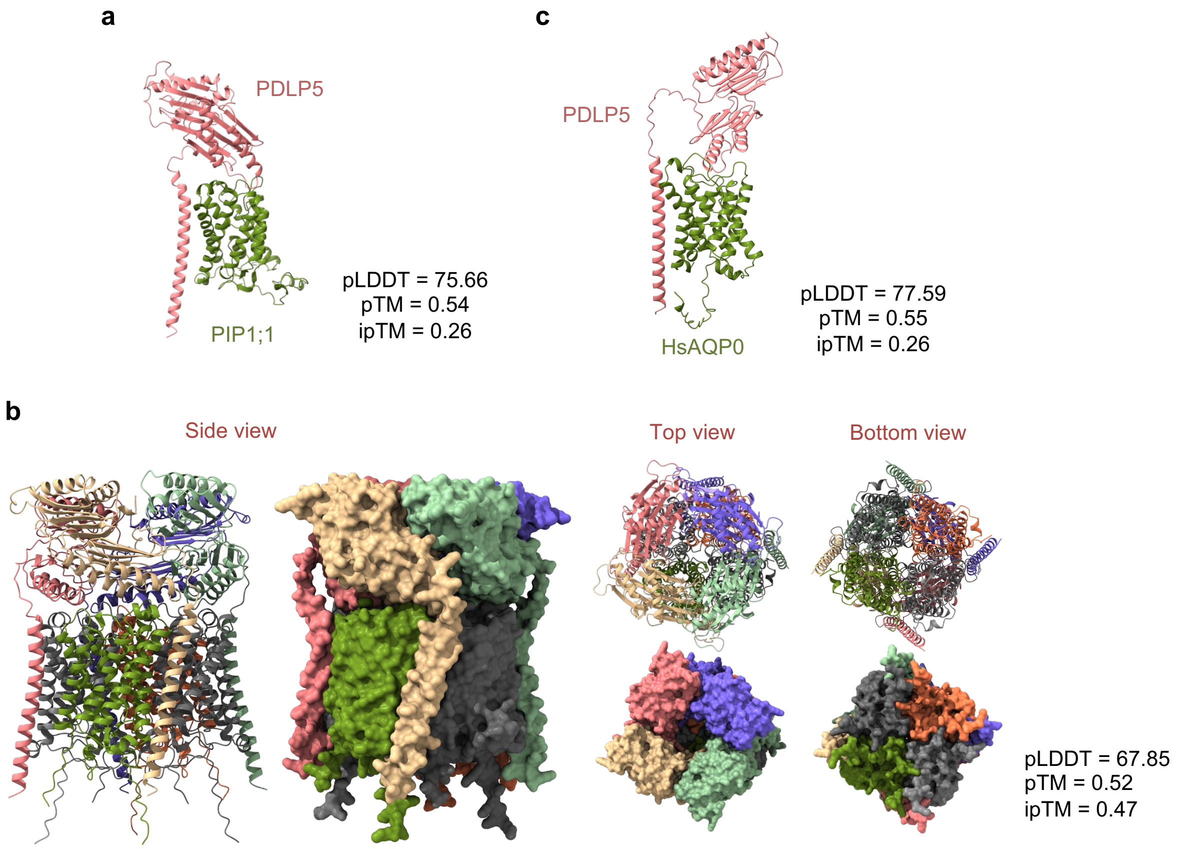


**Extended Data Fig. 1:** AlphaFold predicts PDLP5-AQP complexes. **a,** AlphaFold3 predicts the formation of a PDLP5-PIP1;1 heterodimer; the PDLP5 signal peptide was omitted from the prediction. **b,** AlphaFold3 predicts the formation of a PDLP5-PIP1;1 hetero-octamer. **c**, AlphaFold3 predicts a PDLP5-HsAQP0 heterodimer. Predicted local distance difference test (pLDDT) scores range from 0 (lowest confidence) to 100 (highest confidence), whereas predicted template modeling (pTM) and interfacial predicted template modeling (ipTM) scores range from 0 (lowest confidence) to 1 (highest confidence).
