## Extended Data Fig. 2Extended Data Fig. 1 for "PDLP5 regulates aquaporin-mediated hydrogen peroxide transport in *Arabidopsis*"

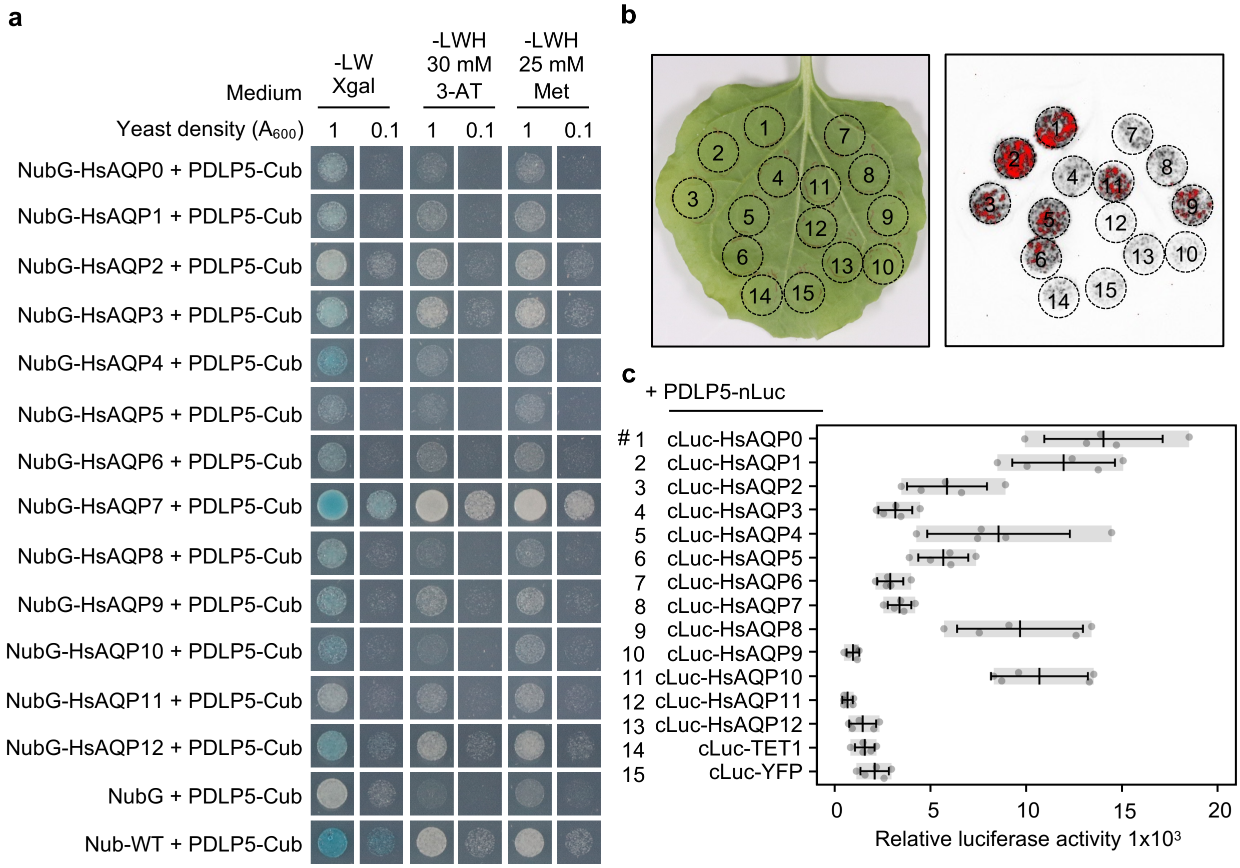


**Extended Data Fig. 2:** PDLP5 physically interacts with human aquaporins (HsAQPs). **a**, Split-ubiquitin yeast two-hybrid assay determines the interaction between PDLP5 and HsAQPs. Nub-WT and NubG were used as positive and negative controls, respectively. To conduct blue/white screening, -LW + X-Gal synthetic dropout plates were utilized, while -LWH synthetic dropout plates were used for histidine auxotrophy selection. To increase the stringency of the assay, 30 mM 3-AT or 25 mM Met was added to the -LWH synthetic dropout plates. **b**, The luciferase complementation assay demonstrates the interaction between PDLP5 and HsAQPs. Agrobacteria carrying the respective plasmids were co-infiltrated into *N. benthamiana* leaves. Two days after infiltration, leaves were sprayed with 1 mM D-luciferin, and luminescence was imaged using an iBright system (top). **c**, Quantification of luminescence was performed using a microplate reader (bottom). Data represent the mean ± SD (n = 5).
