## Extended Data Fig. 3 for "PDLP5 regulates aquaporin-mediated hydrogen peroxide transport in *Arabidopsis*"

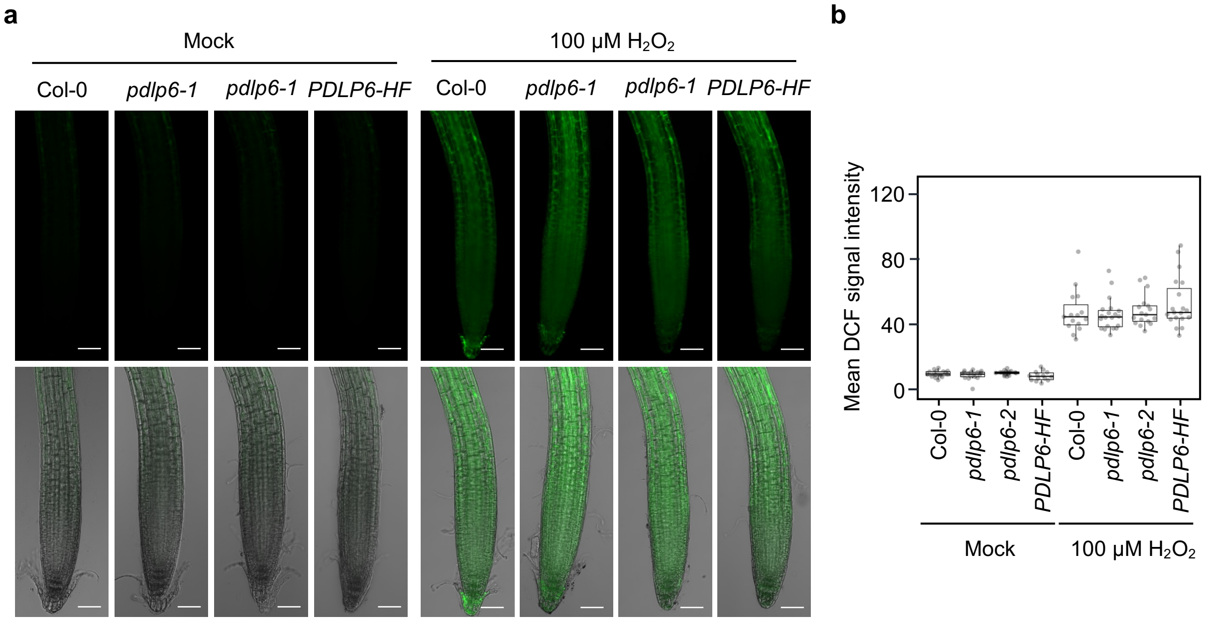


**Extended Data Fig. 3:** PDLP6 does not affect $H_2O_2$ transport in Arabidopsis root tips. **a**, Confocal images show the uptake of $H_2O_2$ in Arabidopsis root tips. 7-day-old Arabidopsis seedlings were incubated with 10 mM $H_2$DCFDA for 15 minutes prior to $H_2O_2$ treatment. DCF signals were imaged 10 minutes after the $H_2O_2$ treatment. Images in the lower panel are merged with bright-field images of root tips. Scale bars = 50 µm. **b,** Quantitative data show the accumulation of $H_2O_2$ in Arabidopsis root tips. The box plot shows the mean with SD. Col-0: n = 17 (mock) and n = 15 (100 µM $H_2O_2$); *pdlp6-1*: n = 17 (mock) and n = 20 (100 µM $H_2O_2$); *pdlp6-2*: n = 17 (mock) and n = 17 (100 µM $H_2O_2$); and *PDLP6-HF*: n = 14 (mock) and n = 19 (100 µM $H_2O_2$).
