## Extended Data Fig. 4 for "PDLP5 regulates aquaporin-mediated hydrogen peroxide transport in *Arabidopsis*"

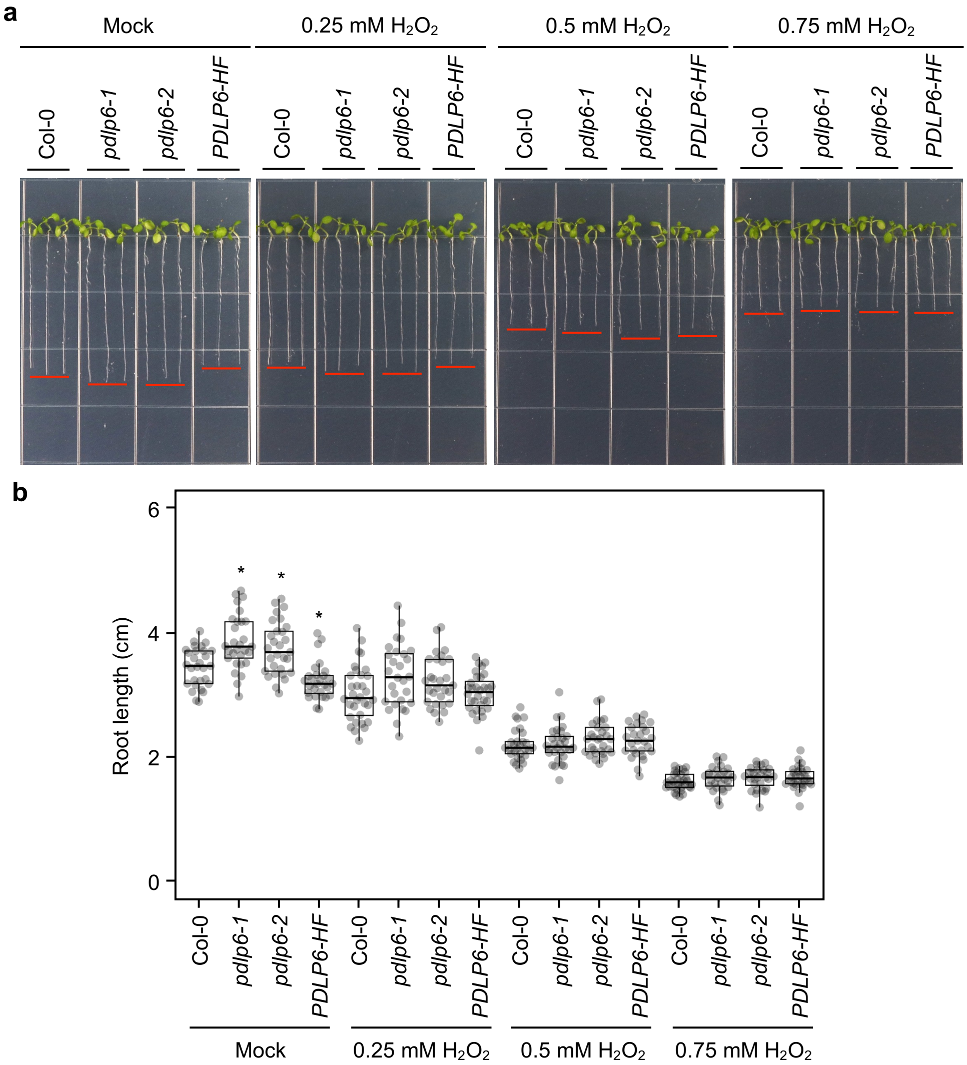


**Extended Data Fig. 4:** PDLP6 does not modulate $H_2O_2$-induced inhibition of root growth in Arabidopsis. **a**, Pictures show root inhibition by $H_2O_2$. Arabidopsis seeds were sown on ½ LS agar plates containing 0.25 mM, 0.5 mM or 0.75 mM $H_2O_2$. Root lengths were measured 10 days after germination. Scale bar = 1 cm. **b,** Quantitative data show root inhibition by $H_2O_2$. The box plot shows the mean with SD. Col-0: n = 29 (mock), n = 32 (0.25 mM $H_2O_2$), n =29 (0.5 mM $H_2O_2$), and n = 32 (0.75 mM $H_2O_2$); *pdlp6-1*: n = 29 (mock), n = 28 (0.25 mM $H_2O_2$), n = 30 (0.5 mM $H_2O_2$), and n = 28 (0.75 mM $H_2O_2$); *pdlp6-2*: n = 29 (mock), n = 29 (0.25 mM $H_2O_2$), n = 29 (0.5 mM $H_2O_2$), and n = 28 (0.75 mM $H_2O_2$); and *PDLP6-HF*: n = 28 (mock), n = 32 (0.25 mM $H_2O_2$), n = 27 (0.5 mM $H_2O_2$), and n = 29 (0.75 mM $H_2O_2$). Asterisks indicate statistically significant differences compared with wild-type Col-0 (*t*-Test; two-tailed; $P < 0.05$).
