## Extended Data Fig. 5 for "PDLP5 regulates aquaporin-mediated hydrogen peroxide transport in *Arabidopsis*"

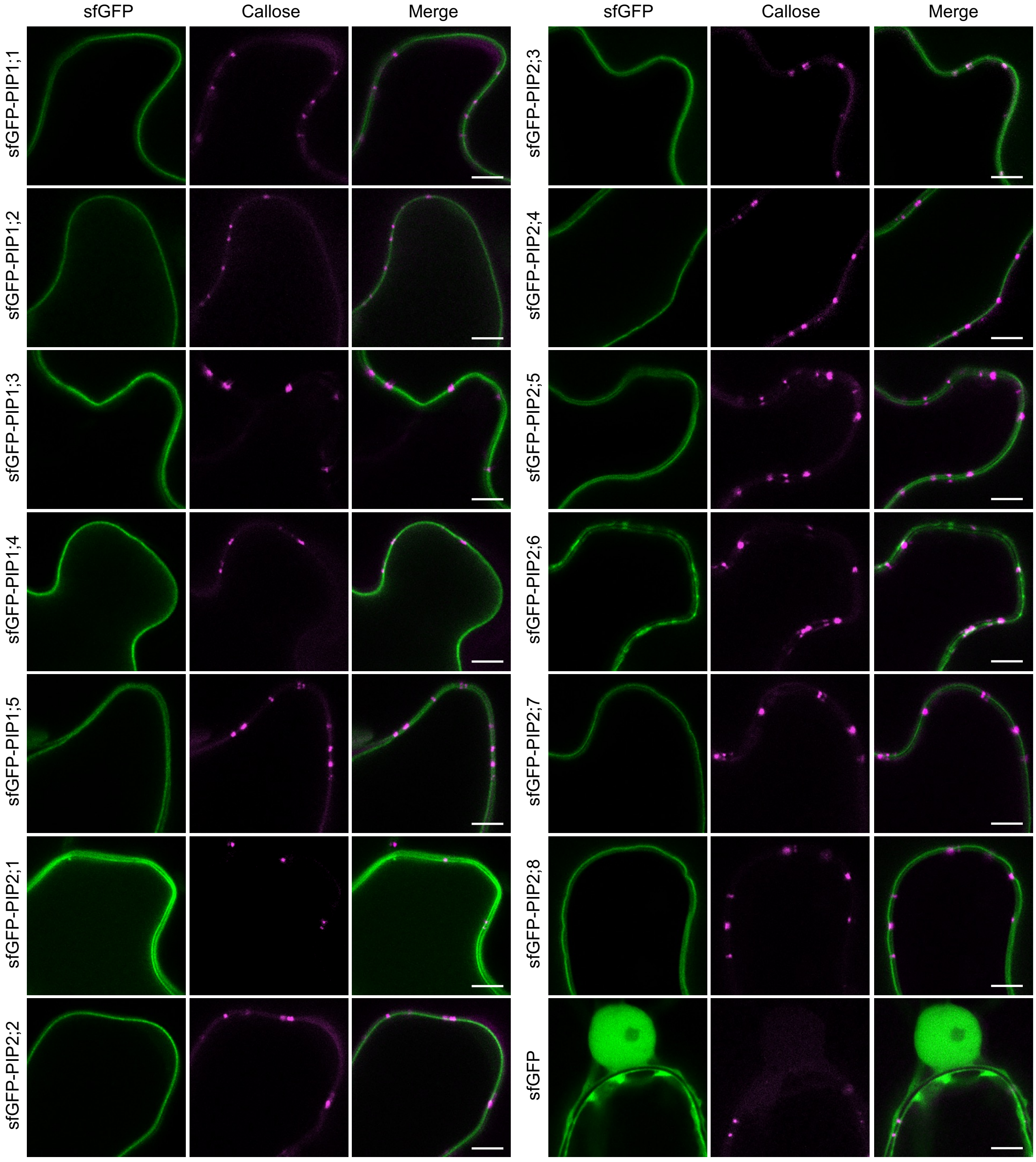


**Extended Data Fig. 5:** Subcellular localization of PIPs. Agrobacteria carrying sfGFP-PIPs were infiltrated into *N. benthamiana* to transiently express the fusion proteins. The subcellular localization of sfGFP fusion protein was detected using confocal microscopy. Aniline blue stains callose at plasmodesmata. A free sfGFP was used as a control to present the expression of the protein in the cytosol and nucleus. Scale bars = 5 µm.
