## Extended Data Fig. 6 for "PDLP5 regulates aquaporin-mediated hydrogen peroxide transport in *Arabidopsis*"

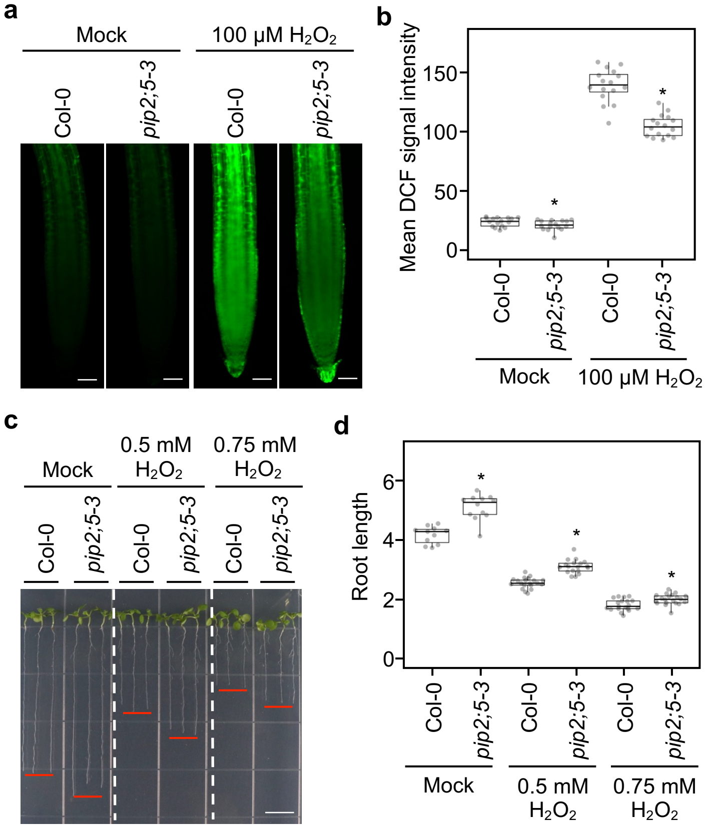


**Extended Data Fig. 6:** PIP2;5 plays a major role in transporting $H_2O_2$ into Arabidopsis root tips. **a,** Confocal images show the uptake of $H_2O_2$ in Arabidopsis root tips. 7-day-old Arabidopsis seedlings were incubated with 10 mM $H_2$DCFDA for 15 minutes prior to $H_2O_2$ treatment. DCF signals were imaged 10 minutes after the $H_2O_2$ treatment. Scale bars = 50 µm. **b**, Quantitative data show the accumulation of $H_2O_2$ in Arabidopsis root tips. The box plot shows the mean with SD. n=16. Asterisks indicate statistically significant differences compared with wild-type Col-0 (*t*-Test; two-tailed; $P < 0.05$). **c**, Pictures show root inhibition by $H_2O_2$. Arabidopsis seeds were sown on ½ LS agar plates containing 0.5 mM or 0.75 mM $H_2O_2$. Root lengths were measured 10 days after germination. Scale bars = 1 cm. **d,** Quantitative data show root inhibition by $H_2O_2$. The box plot shows the mean with SD. Col-0: n = 11 (mock), n = 20 (0.5 mM $H_2O_2$), and n = 19 (0.75 mM $H_2O_2$); *pip2;5-3*: n = 12 (mock), n = 18 (0.5 mM $H_2O_2$), and n = 20 (0.75 mM $H_2O_2$). Asterisks indicate statistically significant differences compared with wild-type Col-0 (*t*-Test; two-tailed; $P < 0.05$).
