## Extended Data Fig. 8 for "PDLP5 regulates aquaporin-mediated hydrogen peroxide transport in *Arabidopsis*"

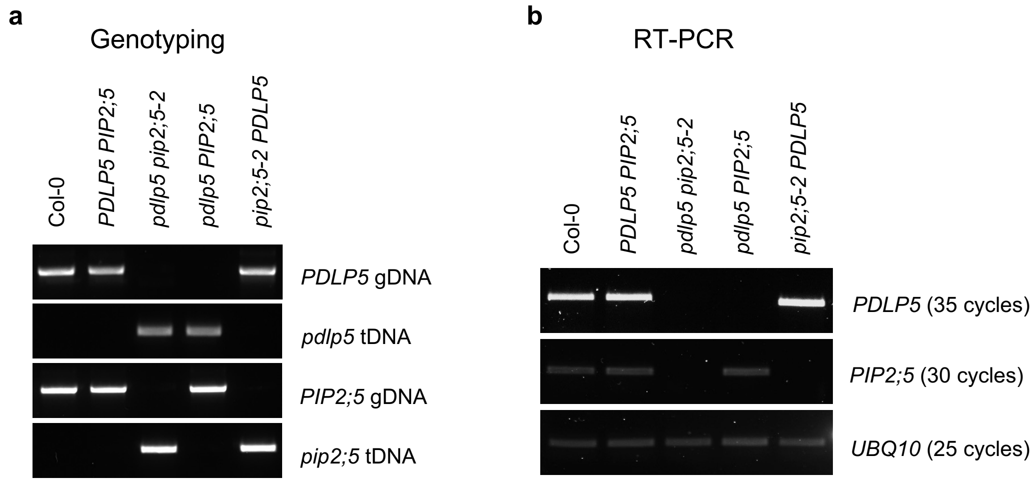


**Extended Data Fig. 8:** Characterization of the *pdlp5 pip2;5-2* double mutant. A genetic cross was performed between *pdlp5* and *pip2;5-2* mutants. The $F_2$ population was used to isolate the following genotypes using a PCR-based genotyping approach: wild type (*PDLP5 PIP2;5*), double mutant (*pdlp5 pip2;5-2*), and single mutants (*pdlp5 PIP2;5* and *pip2;5-2 PDLP5*). **a**, Genomic DNA (gDNA) was amplified using primers flanking the T-DNA insertion site to detect wild-type alleles. T-DNA-specific primers were used to confirm the presence of the T-DNA insertion within *PDLP5* or *PIP2;5*. **b,** RT-PCR was performed to assess the transcript levels of *PDLP5* and *PIP2;5* in wild type, double mutant, and single mutants. *UBQ10* was used as an internal control. The number of PCR cycles used for amplification is indicated.
